# The Thermal Grill Elicits Central Sensitization

**DOI:** 10.1101/2025.02.12.637900

**Authors:** Matthew A. Cormie, David A. Seminowicz, Massieh Moayedi

## Abstract

The thermal grill, in which innocuous warm and cool stimuli are interlaced, can produce a paradoxical burning pain sensation—the thermal grill illusion (TGI). While the mechanisms underlying TGI remain unclear, prominent theories point to spinal dorsal horn integration of innocuous thermal inputs to elicit pain. It remains unknown whether the TGI activates peripheral nociceptors, or solely thermosensitive afferents integrated within the spinal cord. Different types of sensitization have established mechanisms and can inform TGI mechanisms: if the TGI elicits (1) primary hyperalgesia, peripheral nociceptors are likely be activated; (2) secondary hyperalgesia without primary hyperalgesia, spinal integration is likely required; and (3) brush allodynia, wide-dynamic range neurons could be involved in mediating the TGI. Here, we determine whether the TGI elicits primary hyperalgesia, secondary hyperalgesia or brush allodynia. Fifty-two participants underwent individually calibrated phasic thermal grill stimulation. We found that the calibrated TGI elicited primary hyperalgesia, but only in participants with component temperatures in the noxious range (<19°C and >41°C). The TGI also elicited secondary hyperalgesia, even in participants with strictly innocuous thermal inputs. No participants developed brush allodynia. We observed sex differences in primary hyperalgesia: only males exhibited thermal grill-induced primary hyperalgesia. These findings suggest that the TGI is integrated in the spinal dorsal horn, potentially mediated by heat-pinch-cold (HPC) neurons, and, to some degree, by primary nociceptive afferents in males. This study shows that the calibrated TGI may have sex-dependent mechanisms and determines that HPC cells are involved in the illusory sensation of pain from innocuous thermal inputs.

## Introduction

The thermal grill illusion (TGI) is a paradoxical sensory phenomenon causing a perception of pain and burning heat in response to alternating warm and cool innocuous stimuli [5,38,87,94,95]. A key feature of this illusion is the perception of synthetic heat, meaning the overall temperature perception is hotter than either stimulus (warm or cool) in isolation [38]. This experience of heat pain with the TGI has been frequently compared to the burning sensation commonly associated with cold pain [8]. Between 20-75% of individuals report the TGI as painful, showing substantial variability across studies [3,16,58,80,81].

The mechanisms driving the illusory perception of pain in TGI remain a source of debate. Peripherally, the TGI is characterized by its slow and delayed onset [58], and the sensation’s quality is unaffected by ischemic block, which suggests mediation by C-fibres [44]. Two prevailing theories on the mechanism behind the illusion differ in their proposed methods of central integration: the addition theory and disinhibition theory [38,87]. The addition theory posits the TGI is induced by the linear summation of convergent warm and cool input to second-order wide dynamic range (WDR) neurons in the dorsal horn of the spinal cord [48,49]. In contrast, the disinhibition theory posits that TGI is caused by the central disinhibition of second-order heat-pinch-cold (HPC) neurons. Normally, HPC neurons are inhibited by second-order neurons that encode cool stimuli (COOL). However, when a warm stimulus is introduced, the warm primary afferent inhibits the COOL fibres, thus disinhibiting HPC neurons. HPC neurons primarily receive input from C-polymodal nociceptors (CPNs) in the periphery [44], which also encode cool stimuli below the thermoneutral zone (∼24°C) [12,13,29,31–33]. Further, nociceptive specific (NS) neurons showed no change in activity in response to a thermal grill [31]. The TGI is primarily thought to be caused by spinal or supraspinal interactions [16,31,33,39,40,42,48,49,68,72], but it remains unknown whether innocuous warm and cool stimulations activate nociceptive primary afferents [87]—as proposed in the disinhibition theory involving CPNs [31].

To determine whether TGI activates primary nociceptors, or whether the integration is wholly in the CNS can be determined through the investigation of sensitized states. Specifically, primary hyperalgesia would indicate primary nociceptor activation; whereas secondary hyperalgesia, in the absence of primary hyperalgesia, would indicate central integration [57,100]. Furthermore, the second-order neuron encoding the TGI can be delineated through investigation of brush allodynia (also known as stroking hyperalgesia) which is mediated by WDR cells when evoked by noxious thermal heat [24,77].

Therefore, the aims of this study are to determine whether the thermal grill elicits primary, secondary and/or stroking hyperalgesia. Based on extant theories and evidence, we hypothesize that there will be secondary hyperalgesia, but not primary hyperalgesia. We are agnostic of whether brush allodynia can be elicited by the thermal grill, which would determine whether WDRs are involved. This study is the first to use altered states to disentangle the mechanisms of the TGI.

## Methods

### Participants

Sixty-seven participants between the ages of 18 and 45 completed the study approved by the University of Toronto Research Ethics Board (Protocol#: 45598). Exclusion criteria for the study were (1) current or history of chronic pain; (2) major neurological or psychiatric diseases; (3) vascular diseases; (4) skin diseases (i.e. psoriasis or open sores); (5) self-reported opioid use; (6) daily use of medication known to affect pain sensitivity (e.g., NSAIDS, anxiolytic, antispastic, acetaminophen); (7) a score ≥20 on the Beck’s Depression Inventory II (BDI-II) [10]; (8) indication of suicidal ideation (>1 on item 9 of the BDI-II); (9) a score of <28 on Mini Mental State Exam (MMSE; see *Experimental Design*, below, for rationale) [41]; and (10) average pain intensity ratings <5/100 to thermal grill stimulation during TGI calibration (described in detail in the *Thermal Grill Calibration* section). Of the 67 participants, 15 were excluded due to being a TGI “non-responder” (average TGI calibration pain intensity <5/100; n=14) or scored >20 on the BDI-II (n=1). Fifty-two participants (26 females, 26 males; mean age ± SD: 23.5 ± 4.8 years) were invited to undergo the full experiment. To explore whether phasic thermal grill stimulation could induce secondary hyperalgesia, a pilot study (n = 5) was conducted. A one-sample t-test was used to determine whether the area of secondary hyperalgesia was significantly greater than 0 cm². Based on the observed effect size (Cohen’s *d* = 1.444), a power analysis with a desired power of 0.95 indicated that a minimum of 7 participants would be needed to replicate the effect, or 14 participants to allow for adequately powered sex-disaggregated analyses (i.e., two groups of 7 participants). The final sample size (n = 52) was purposefully oversampled to ensure reliable and generalizable results [55].

### Experimental Design

Participants underwent a single behavioural session lasting one hour (Figure 1). After consenting to procedures, participants were first screened for moderate depression with the BDI-II [10] and cognitive impairment with the MMSE [41], as per the exclusions above. Cognitive impairment was assessed to ensure participants’ alertness and were able to understand and follow the experimenter’s directives. Next, they completed questionnaires for measures known to be associated with pain sensitivity.

**Figure 1:**
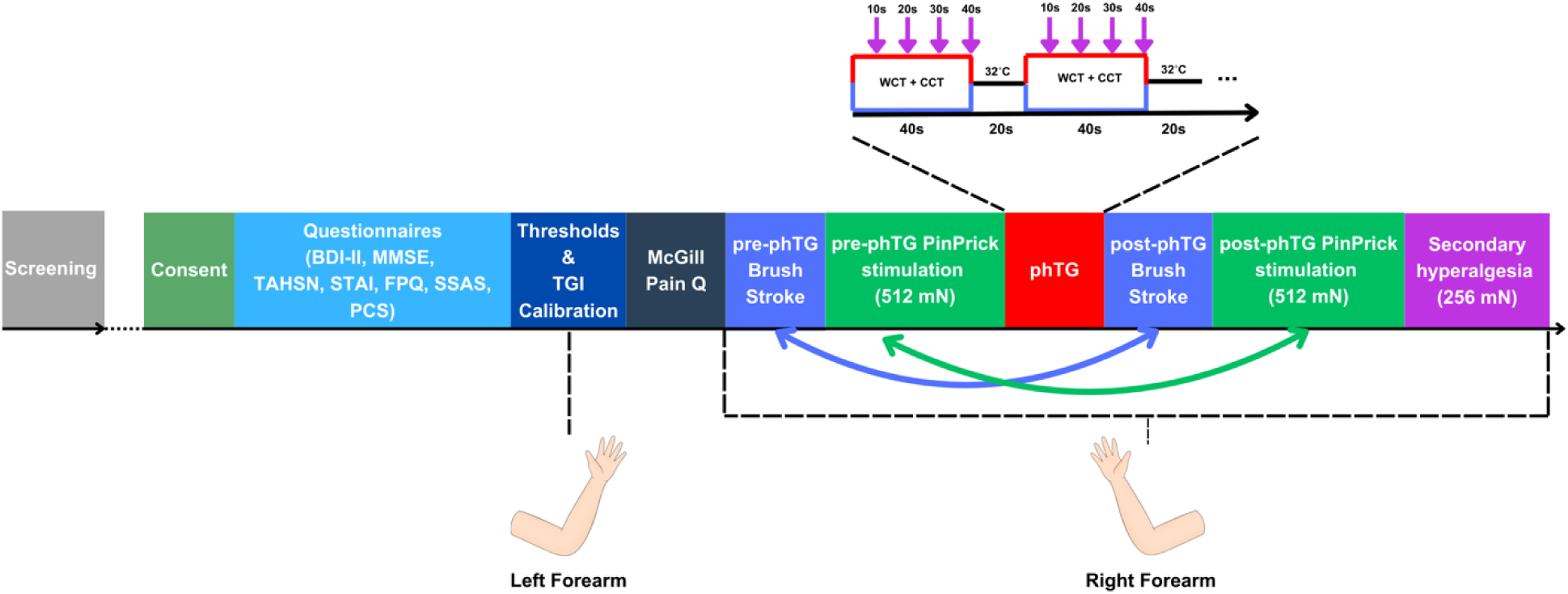
Timeline of study visit. Curved arrows represent behavioural measures repeated pre-phTG and post-phTG. Thermal pain threshold and TGI calibration occurred on the left forearm, all other stimuli were on the right. The inset provides a schematic of the phTG stimulation paradigm, consisting of 40s stimulation and 20s interstimulus interval, repeated eight times. Purple arrows indicate when participants are prompted to provide ratings.

Note that the thermal grill comprises a warm component and a cool component, each of which must be perceived as non-painful. Thus, we measured participants cold pain and heat pain thresholds (CPT and HPT, respectively) to determine the non-painful range for component temperatures of the thermal grill. These procedures were performed on the left volar forearm, with a two-minute break between the CPT and HPT. The HPT was followed by another two-minute break, after which thermal grill component temperatures were calibrated on participants’ left volar forearm. We then assessed whether participants were TGI responders. TGI non-responders’ sessions ended at this point. TGI responders were asked to provide qualitative descriptors of the TGI sensation using the McGill Pain Questionnaire [70]. Next, participants underwent baseline sensitization measures at the stimulation site on their right volar forearm.

Participants then underwent a modified phasic heat pain paradigm [59,78]; instead of receiving noxious heat stimuli, we performed a phasic thermal grill (phTG) stimulation with individually calibrated component temperatures. Finally, participants underwent post-phTG sensitization measures. This concluded the study session. Participants were financially compensated for their time.

#### Questionnaires

*Beck’s Depression Inventory-II (BDI-II)* is a questionnaire that measures participants’ depression levels. The questionnaire consists of 21 items, scored on a 4-point Likert scale ranging from 0-3. Scores can range from 0-63 [10].

The *Mini Mental State Examination (MMSE)* is a questionnaire that assesses participants’ alertness and cognitive impairments. The exam consists of 19 questions asking participants to follow the experimenter’s directives. Scores can range from 0-30 [41].

*State-Trait Anxiety Inventory (STAI*) is a questionnaire that measures participants’ state and trait anxiety. The first 20 questions ask participants questions to determine their anxiety level at the current moment (state anxiety). The following 20 questions are used to determine the participants general level of anxiety (trait anxiety). State and trait anxiety levels are computed separately. Questionnaire scores range from 20-80, for each subscale [90].

The *Pain Catastrophizing Scale (PCS)* measures participants’ emotions and thoughts when experiencing pain. Catastrophizing score is calculated on a 4-point Likert scale across 13 items. The questionnaire has three validated subscales measuring magnification, rumination, and helplessness when experiencing pain. Scores can range from 0-52 [92].

The *Fear of Pain Questionnaire-III (FPQ-III)* measures participants’ fears towards painful experiences. The questionnaire consists of 30 items, scored on a 5-point Likert scale ranging from 1 (not at all) to 5 (extreme). Scores can range from 30-150 [69].

The *Somatosensory Amplification Scale (SSAS)* measures participants’ tendency to detect and focus on physiological states and uncomfortable bodily sensations (i.e. heart rate). The questionnaire consists of 10 items, scored on a 5-point Likert scale ranging from 1 (strongly disagree) to 5 (strongly agree). Scores can range from 10-50 [9].

The *Toronto Academic Health Science Network (TAHSN)* demographics questionnaire asks participants to self-report personal demographics information: age, sex, gender, race, sexual orientation, spiritual or religious affiliation, disability status, and family income.

#### Thermal Stimulation Paradigm

For all thermal stimulations (HPT, CPT, CCT, WCT, and TGI), we used the TCS II thermal stimulating device (QST.Lab, Strasbourg, France) with a T11 thermode. This thermode has five separate “grills” that can be independently controlled and set to different temperatures (see Figure 2). We only used 4 of the 5 grills to have a stimulation area (7.4 mm x 24.2 mm = 178.08 mm^2^ per grill, or ∼7.16 cm^2^ total)), and to ensure equal number of grills for cool and warm. The fifth grill was set to the fixed baseline temperature (32°C). For thermal grill stimulations, grill 1 is set to baseline, grills 2 and 4 to the calibrated warm component temperature (WCT), and grills 3 and 5 to the calibrated cool component temperature (CCT). The thermode is placed on top of the volar forearm and held in place by the experimenter for all stimulations, applying no additional downward pressure beyond the gravitational force on the thermode. Throughout the study, ‘baseline’ refers to a thermode temperature of 32°C.

**Figure 2:**
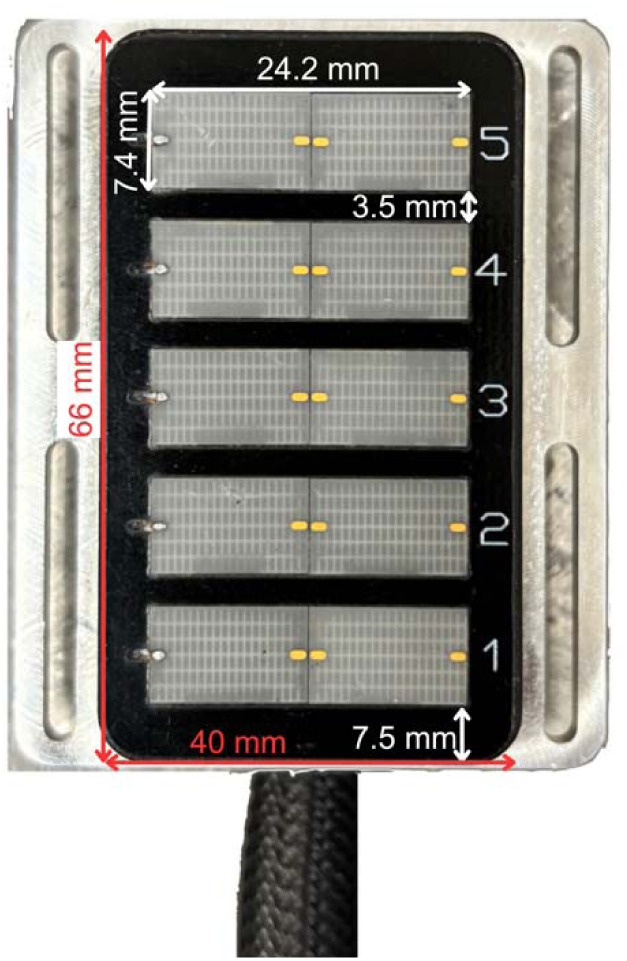
QST.Lab T11 thermode. The T11 thermode consists of five independently controlled stimulation surfaces (‘grill’). Each grill is 7.4 mm x 24.2 mm, for an area of 179.08 mm. There is a 3.5 mm spacing between the grills. The total stimulation surface of the four grills utilized in the study is 716.32 mm2. The total probe dimensions (red) are 66 mm x 40 mm, for an area of 26.4 cm2 in contact with the participant.

#### Pain Threshold Calibration

The HPT and CPT calibration were performed as previously done by Mitchell et al. [72]. Specifically, HPT and CPT were determined on the left volar forearm of participants using a staircase method (method of levels). We first determined CPT, followed by a 2-minute break, after which we determined HPT.

The volar forearm was divided into 3 non-overlapping regions, with the most proximal stimulus being 4 cm from the elbow, and the most distal being 4 cm from the wrist. The spacing between each is variable depending on the length of the participant’s forearm.

For CPT calibration, the first stimulus is in the thermoneutral zone at 25°C. Each stimulus lasted 5 seconds, including ramp up/down at 25°C/second. After every stimulus, participants were prompted to verbally rate whether the stimulus was painful (binary “pain” / “no pain”). The thermode was then moved to another stimulation region on the volar forearm. If the participant reported no pain, the stimulus was decreased by 5°C. If the participant reported pain, the stimulu was increased by 1°C. The interstimulus interval was ≥5 seconds, depending on the speed at which participants provided ratings. The procedure was continued until the participant rated a temperature as painful over 3 consecutive trials. There was no maximum number of stimuli given to participants, but no participants exceed 15 stimuli. Therefore, participants are given ample time to feel the stimulation and provide a response. The minimum temperature given was 0°C.

For HPT calibration, the same procedure was repeated but the first stimulus was 35°C and was increased by 5°C intervals until pain was reported, then decreased by 1°C intervals. Again, the procedure was continued until the participant rated a temperature as painful over 3 consecutive trials. The maximum temperature given was 45°C.

#### Thermal Grill Calibration

Studies have found that 20-75% of individuals find the TGI painful [3,16,58,80,81]; therefore, to explore the individual differences in the pain percept associated with the illusion, participants received individually calibrated thermal grill component temperatures. The thermal grill calibration occurred after CPT/HPT calibration and was used to find WCT and CCT that were consistently reported as non-painful by participants. For the CCT, stimulation began 1°C above the participant’s CPT, if the participants reported pain, the stimulus was increased 1°C repeatedly until a stimulation temperature that is not painful over three trials was identified. Again, to avoid sensitization, the thermode was moved between stimulations. Then following a 2-minute break, the same paradigm was repeated to determine WCT, with the first stimulus 1°C below participants HPT, and repeatedly reduced by 1°C. The minimum CCT used was 5°C and the maximum WCT used was 42°C. After determining participants WCT and CCT, participants were given another 2-minute break before receiving three thermal grill stimulations, to ensure they report the thermal grill as painful. Participants were prompted to provide pain intensity and unpleasantness ratings after each stimulation. Pain intensity ratings were on a verbal numerical rating scale (vNRS) with anchors 0 = “no pain at all” and 100 = “the worst imaginable pain”.

Stimulus unpleasantness ratings were also on a vNRS but with anchors 0 = “not unpleasant at all” and 100 = “the most unpleasant sensation imaginable”. These rating scales were used throughout the study, specifically during phTG stimulation and for mechanical punctate stimulation.

There is no consensus on criteria to determine TGI responders and non-responders. We have a employed a modified version of TGI calibration, as described by Mitchell et al. [72], to capture the individual variability in pain percepts and responsiveness to the TGI. If participants provided a mean pain intensity rating less than 5/100 across the three trials, the study session was terminated, and the participant deemed a TGI non-responder. The threshold of 5/100 was selected to ensure the stimulations were reliably painful, but not overly strict as to exclude mild TGI responders. Finally, after determining TGI responsiveness, participants are given one final thermal grill stimulation and prompted to select descriptor words from the MPQ to describe the TGI sensation. Within the MPQ, descriptors are grouped together under word sets. If a participant selected multiple descriptors within the word set, the most intense word was scored.

#### Phasic Thermal Grill (phTG) Stimulation Paradigm

To elicit sensitization, participants received a phTG stimulation on their right volar forearm. The phTG occurred after CPT/HPT calibration, TGI component temperature calibration, and baseline brush and mechanical pinprick measures. The phTG stimulation consisted of eight 40 s calibrated thermal grill stimuli, with a 20 s inter-stimulus interval. As the intent behind the phasic stimulation was to induce sensitization, the thermode was not moved between the eight phTG stimuli and was maintained within a designated stimulation site on the volar forearm (see Figure 3). Participants verbally provided both pain intensity and unpleasantness ratings every 10 s during stimulation, resulting in four ratings per stimulation across eight stimulations, for 32 ratings on each rating scale (32 for pain intensity, 32 for stimulus unpleasantness). Throughout the paradigm participants are prompted to rate their pain intensity and stimulus unpleasantness at every 10s (four times during each 40s phasic stimulation, a total of 32 ratings for the phTG paradigm).

**Figure 3:**
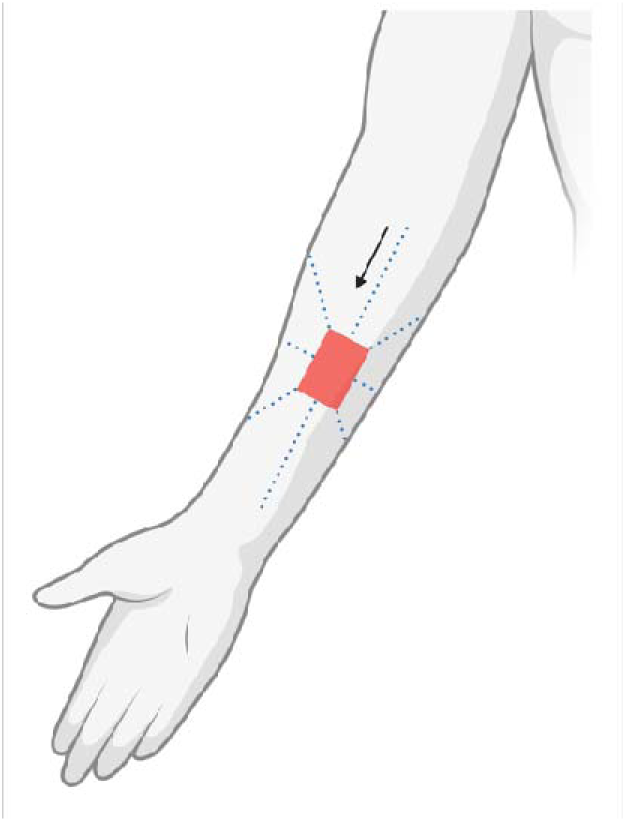
Secondary hyperalgesia map. The map consists of 8 arms radiating outwards from the stimulation site, with each arm containing 14 dots at 0.5cm increments (total distance in each direction = 7cm). Stimulation begins at the point farthest from the stimulation site, and moved inwards until the edge of the stimulation site (direction in arrow) is reached or the participant reports and increase in intensity of the punctate stimulus. This is repeated for all 8 radial arms. Created in BioRender. Moayedi, M. (2025) https://BioRender.com/d8bc9lh.

#### Innocuous Brush Stimulus (Brush Allodynia)

To determine whether phTG stimulation can induce brush allodynia, participants received calibrated brush stimulations (SENSELab Brush 05, Somedic SENSELab AB, Sösdala, Sweden) before (pre-phTG) and after (post-phTG) they complete phTG paradigm, within the stimulation site. Both pre- and post-phTG, participants received three brush strokes and were prompted to give a binary response about whether they felt pain to the prompt: “Is it painful, yes or no?” In total, participants provided six pain ratings for brush allodynia (three pre-phTG, three post-phTG). We operationalized brush allodynia as pain to innocuous brush stimulus, within the area of stimulation, post-phTG. The pre-phTG brush stimulations were to ensure any pain reported post-phTG to the brush stimulus was indeed allodynic.

#### Primary Mechanical Hyperalgesia

To determine primary mechanical hyperalgesia, participants receive a mechanical punctate pinprick with a 512mN probe (PinPrick stimulator, MRC Systems GmbH, Heidelberg, Germany) pre- and post-phTG stimulation, within the stimulation site. Pre-phTG, participants receive three punctate pinprick stimuli over the stimulation site of grills 3, 4, and 5. Participants are prompted to rate their pain intensity using the vNRS after each stimulation. Post-phTG, participants undergo the same procedure. Therefore, participants reported pain intensity ratings at two time points: pre-phTG and after the entire phTG (post-phTG). In total participants provided six pain intensity ratings (three pre-phTG, three post-phTG). The magnitude of primary mechanical hyperalgesia was calculated as the difference between means of post-phTG pain intensity ratings and pre-phTG pain intensity ratings. We operationalized primary mechanical hyperalgesia as a significant increase in average pain intensity to mechanical pinpricks post-phTG, compared to pre-phTG.

#### Secondary Hyperalgesia

We operationalized secondary hyperalgesia as an area of increased pain intensity to a noxious stimulus surrounding the area of stimulation, as previously done [1,78]. Pre-phTG, a secondary hyperalgesia map was drawn onto the participant’s right volar forearm (Figure 3). Post-phTG the area of secondary hyperalgesia was determined. A 256mN PinPrick probe is used to stimulate each dot along the eight radial arms starting with the most proximal line and moving counterclockwise. Stimulations begin at the furthest dot and move inwards at a rate of ∼1Hz.

Participants are prompted to notify the researcher when they experience an increase in intensity of the pinprick stimulation, compared to the previous ones of along the same radial arm. The area of secondary hyperalgesia was calculated using ImageJ [84], by taking the area of sensitivity around the stimulation site and subtracting the area of the thermode.

### Statistical Analyses

All frequentist statistical analyses were performed using SPSS Statistics v29.0.2.0 (IBM SPSS Statistics, Armonk, NY, USA) and all data visualization was performed using GraphPad’s Prism v10 (GraphPad Software, Boston, Massachusetts USA). Bayesian statistical analyses were performed using JASP v.0.19.2 (JASP Team, 2024) to supplement frequentist analyses. Bayesian analyses are different from frequentist analyses as they quantify evidence in favour of the null versus alternative hypotheses. Frequentist analyses can only provide a probability that the null hypothesis is rejected (with 95% certainty) [103]. Therefore, Bayesian analyses complement frequentist analyses, especially in cases of null findings, as frequentist statistics cannot accept the null [85]. Bayesian analyses allow for an additional layer of statistical confidence in our conclusions.

In Bayesian statistics, each observation is treated as evidence that updates prior beliefs to produce a posterior probability—the probability assigned to a hypothesis after accounting for both the observed data and prior information. While posterior probabilities provide a direct estimate of belief in a hypothesis, Bayes factors (BFs) are used to compare the relative evidence for two competing hypotheses (typically H_0_ vs. H_1_). BFs can range from 0 to ∞, with set ranges to determine the amount of evidence: 1<BF_10_<3 indicate anecdotal evidence for H_1_; 3<BF_10_<10 indicates moderate evidence for H_1_; 10<BF_10_<30 indicates strong evidence for H_1_, and 30<BF_10_<100 indicates very strong evidence for H_1_; 100<BF_10_ indicates extreme evidence for H_1_. Ranges of evidence for H_0_ are similar but defined as 1/BF_10_ values (e.g., 0.33<BF_10_<0.1 indicates moderate evidence for H_0_). For one-sided tests, BF_0+_ is evidence for the H_0_ for the positive tail, and BF_0-_ is evidence for H_0_ for the negative tail (BF_+0_ and BF_-0_ are the reciprocal of these tests, respectively). All Bayesian analyses were run with 10,000 permutations, where appropriate. For an in-depth of review of Bayesian statistics, see [101].

The Shapiro-Wilk normality test was performed to test whether outcome measures were normally distributed. We investigate whether there are sex differences in sample characteristics and whether phTG induces sensitization, as measured by our three primary outcome measures: brush allodynia, primary mechanical hyperalgesia, and secondary hyperalgesia.

Throughout the results, effect sizes are reported alongside statistical analyses results. For parametric analyses, a Cohen’s *d* effect size was reported. These effect sizes can be interpreted as small (d = 0.2), medium (d = 0.5), and large (d ≥ 0.8) [26,27]. For non-parametric analyses, a rank-biserial correlation (*r*) effect size was reported [11]. These effect sizes can be interpreted as small (r = 0.1), medium (r = 0.3), and large (r ≥ 0.5) [46].

#### Sample characteristics

Analyses were performed for the whole group, as well as in sex-disaggregated groups. Independent samples t-tests, or Mann-Whitney U tests if the variable was not normally distributed, were run to ensure male and female participants did not differ significantly in age, CPT, HPT, WCT, CCT, and questionnaire measures. Significance was set at p<0.05.

#### Brush Allodynia

No statistical test was performed to determine brush allodynia, as no participant reported pain to the brush stroke in the stimulation site pre and post-phTG.

#### Primary Mechanical Hyperalgesia

To test whether the phTG induced primary mechanical hyperalgesia, averaged pre- and post-phTG pain intensity ratings to mechanical punctate stimuli within the stimulation site were compared. The null hypothesis of the analysis was that there is no significant difference in average pain intensity post-phTG compared to pre-phTG. The alternative hypothesis was that average pain intensity ratings post-phTG were significantly greater than pre-phTG and was thus a one-tailed hypothesis. As both pre- and post-phTG pain intensity ratings to the 512mN mechanical pinprick were not normally distributed (p < 0.001), a related-samples Wilcoxon signed rank test was run, with significance set at p < 0.05. A Bayesian Wilcoxon signed rank test was also run to determine the strength of evidence for the null and alternative hypotheses.

To determine whether there were any differences in primary mechanical hyperalgesia within each sex, the same analyses was performed with sex disaggregated samples. The null and alternative hypotheses were identical to whole group sample analyses. We further investigated if there were sex differences in primary hyperalgesia. As the data was not normally distributed, we computed the difference in ratings between post-phTG and pre-phTG (magnitude of primary hyperalgesia, as mentioned previously) and performed a Mann-Whitney U test to compare primary hyperalgesia between groups. Again, to determine the strength of evidence for the null and alternative hypotheses, a Bayesian Mann-Whitney U test was performed. The null hypothesis was there is no significant difference in the magnitude of primary hyperalgesia in males versus females. The alternative hypothesis was the magnitude of primary hyperalgesia was significantly different between males and females and was therefore a two-tailed hypothesis.

#### Secondary Hyperalgesia

To test whether phTG induced secondary hyperalgesia, a one-sample t-test was run comparing the area of secondary hyperalgesia in each individual with a test value set to 0 cm^2^. A Bayesian one-sample t-test was again run to determine strength of evidence for the hypotheses. The null hypothesis was the area of secondary hyperalgesia was not significantly different to the test value of 0 cm^2^, while the alternative hypothesis was the area of secondary hyperalgesia was significantly greater than 0 cm^2^ and thus was a one-tailed hypothesis.

To determine whether there were any differences in secondary hyperalgesia within each sex, the same analyses was performed with sex disaggregated samples, with the same null and alternative hypotheses. We also performed independent samples t-test was run to compare male and female participants’ area of secondary hyperalgesia, with significance set at p<0.05. Again, a Bayesian independent samples t-test was performed to determine the strength of evidence in favour of the hypotheses. The null hypothesis was there is no significant difference in the area of secondary hyperalgesia in males versus females, while the alternative was that there is a significant difference and thus was a two-tailed hypothesis.

#### Contribution of WCT and CCT to primary outcome measures

To determine whether component temperatures affected primary outcome measures, Bayesian ANCOVAs were run with participants’ WCT and CCT as covariates. For primary hyperalgesia, a repeated measures Bayesian ANCOVA was performed. For both, the null hypothesis was WCT and CCT do not significantly contribute to the model, while the alternative was that they do significantly contribute and thus was a two-tailed hypothesis.

#### Analysis of sensitization in a subsample with WCT and CCT in the innocuous range

The thermal noxious range varies significantly across studies—depending on stimulation parameters (location, intensity, size, ramp rate, and duration of the stimulus), the species being tested, as well as the nociceptor type stimulated [17,50,96,98,99]. Based on the activation thresholds of transient receptor potential (TRP) villanoid 1 (TRPV1) of 42°C, which transduce noxious heat [20,56,73], and TRP ankyrin 1 (TRPA1) of 17°C, which transduce noxious cold [60,62,79,91], the innocuous range in humans is largely thought to be between 17°C and 42°C. In the present study, some participants received component temperatures putatively in the noxious range, even if they perceived the stimuli as non-painful. We wanted to determine whether participants who received strictly innocuous stimuli from the thermal grill developed primary mechanical and/or secondary hyperalgesia. As the classical thermal grill (40°C and 20°C) is largely considered innocuous [3,6,16,38,72,87], we operationalized a conservative innocuous range of < 41°C and > 19°C. Selecting only participants whose WCT was < 41°C and whose CCT was > 19°C resulted in a subsample of n = 7 (2 males, 5 females). All measures were normally distributed (p > 0.05).

For primary hyperalgesia, we performed a paired t-test in this subgroup, with significance set at p < 0.05. Bayesian paired t-tests were also run to determine the strength of the evidence in support of the null and alternative hypotheses. For secondary hyperalgesia, we performed a one-sample t-test in this subgroup, with significance set at p<0.05. Bayesian one sample t-tests was also run to determine the strength of evidence for the null and alternative hypotheses. For both primary and secondary hyperalgesia analyses, null and alternative hypotheses were the same as the whole sample analyses.

#### Associations between TG-induced sensitization and psychosocial measures

The relationship between psychosocial measures and the TGI percept remain understudied. Studies in clinical populations have found that the TGI is less painful in those with major depressive disorder [14]. In healthy populations rumination and interoceptive accuracy were associated with TGI pain responses [82], while the induction of a sad mood exacerbates TGI pain [15]. Psychosocial factors such as anxiety, pain catastrophizing, fear of pain and somatosensory amplification have previously been shown to affect pain sensitivity [54,69,71,76,92]. To further explore the influence of psychosocial factors in the TGI, exploratory Bayesian correlation analyses were performed to determine whether there was evidence to support the hypothesis that psychosocial measures (BDI-II, STAI State, STAI Trait, PCS, FPQ, and SSAS) were related to TGI pain intensity ratings (average rating during phTG), magnitude of primary hyperalgesia, and secondary hyperalgesia. Further, as all measures except for secondary hyperalgesia were not normally distributed (p<0.05), Kendall’s tau correlations were performed. The null hypothesis was that there was no correlation between psychosocial measures and outcome measures (pain intensity, magnitude of primary hyperalgesia, and secondary hyperalgesia), while the alternative was that there was a significant correlation (two-tailed).

#### Sex Disaggregated Analyses

To ensure any observed difference in sensitization metrics were not driven by other physiological or psychosocial factor, independent sample t-tests were run to determine if there were any sex differences in HPT, CPT, WCT, CCT, average pain intensity during phTG, STAI state, STAI Trait, PCS, FPQ, and SSAS. If values for either sex was not normally distributed, non-parametric Mann-Whitney U tests were performed. To supplement these analyses, complimentary Bayesian independent t test/Mann-Whitney U tests were performed to determine the evidence for the null and alternative hypotheses. For these analyses, the null hypothesis was there is no significant difference between males and females, while the alternative hypothesis was there was a statistical difference (two-tailed).

## Results

### Sample Demographics and Questionnaire Data

Fifty-two sex-matched participants (26 females, 26 males; age ±SD: 23.5 ±4.8 years) were included in the final analysis. Table 1 provides a summary of demographic information of the sample, including self-reported gender, sexual orientation, race, and religious or spiritual affiliation. Data from participants psychological and behavioural questionnaire data is summarized in Table 2.

**Table 1:**
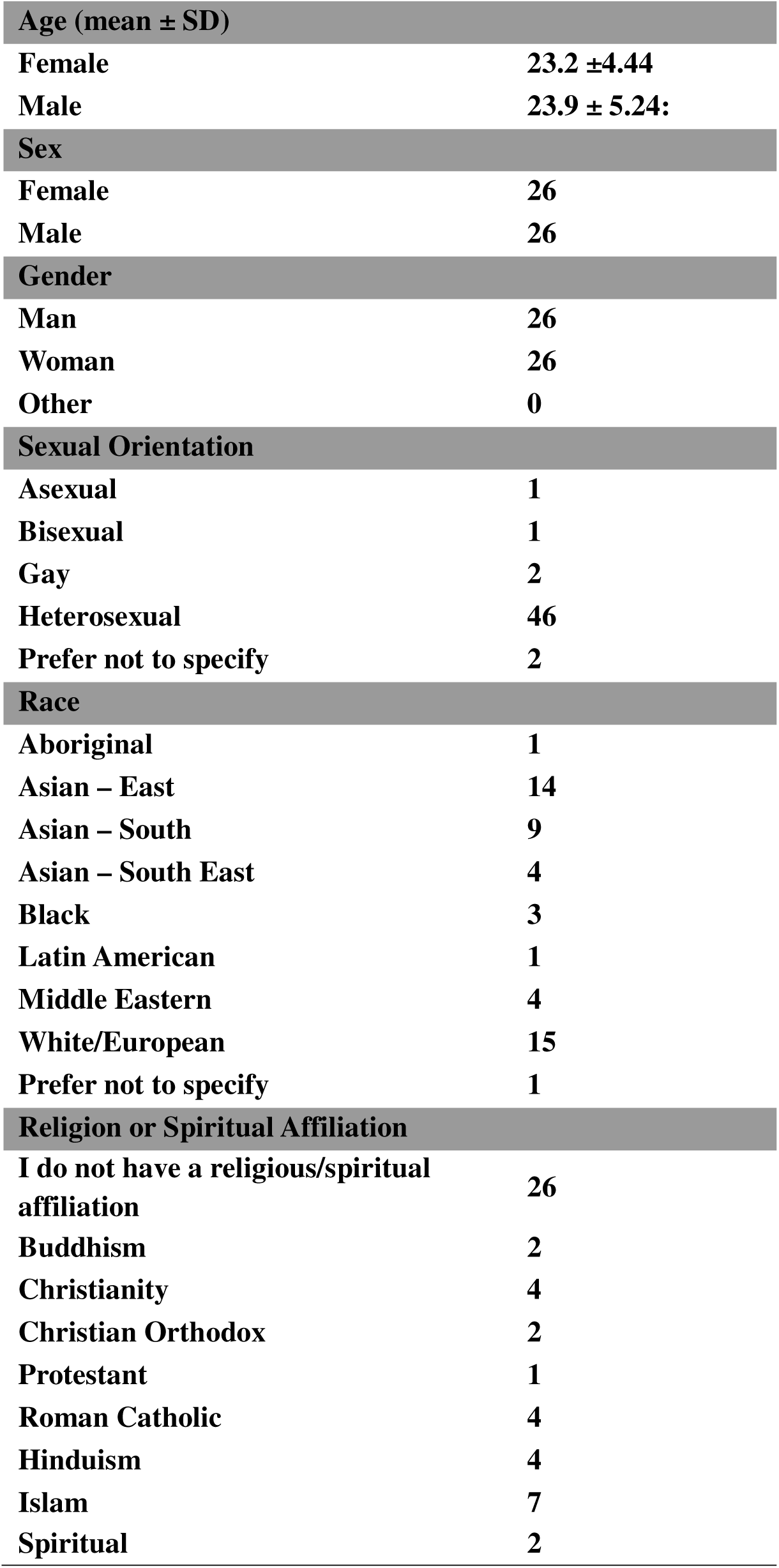
Demographic information of the study sample.

**Table 2:**
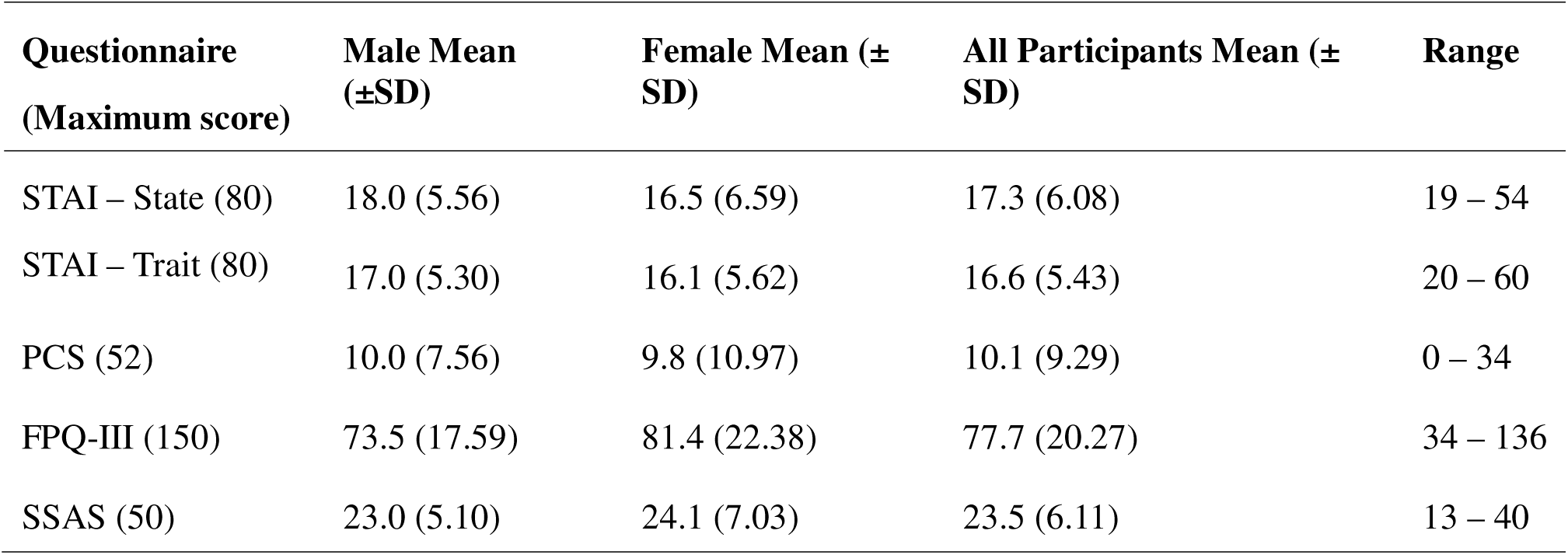
Sample psychosocial questionnaire measures. Group and sex-specific summaries of psychosocial questionnaire data, including means and ranges.

### Calibrated Thermal Grill Component Temperatures

Participants underwent CPT and HPT calibration using a method of levels to determine their thermal pain ranges. Participants had a median [IQR range] HPT of 43.5°C [42.5 – 44.0] (Figure 4A), and CPT of 2.0°C [0.0 – 11.0] (Figure 4B).

**Figure 4:**
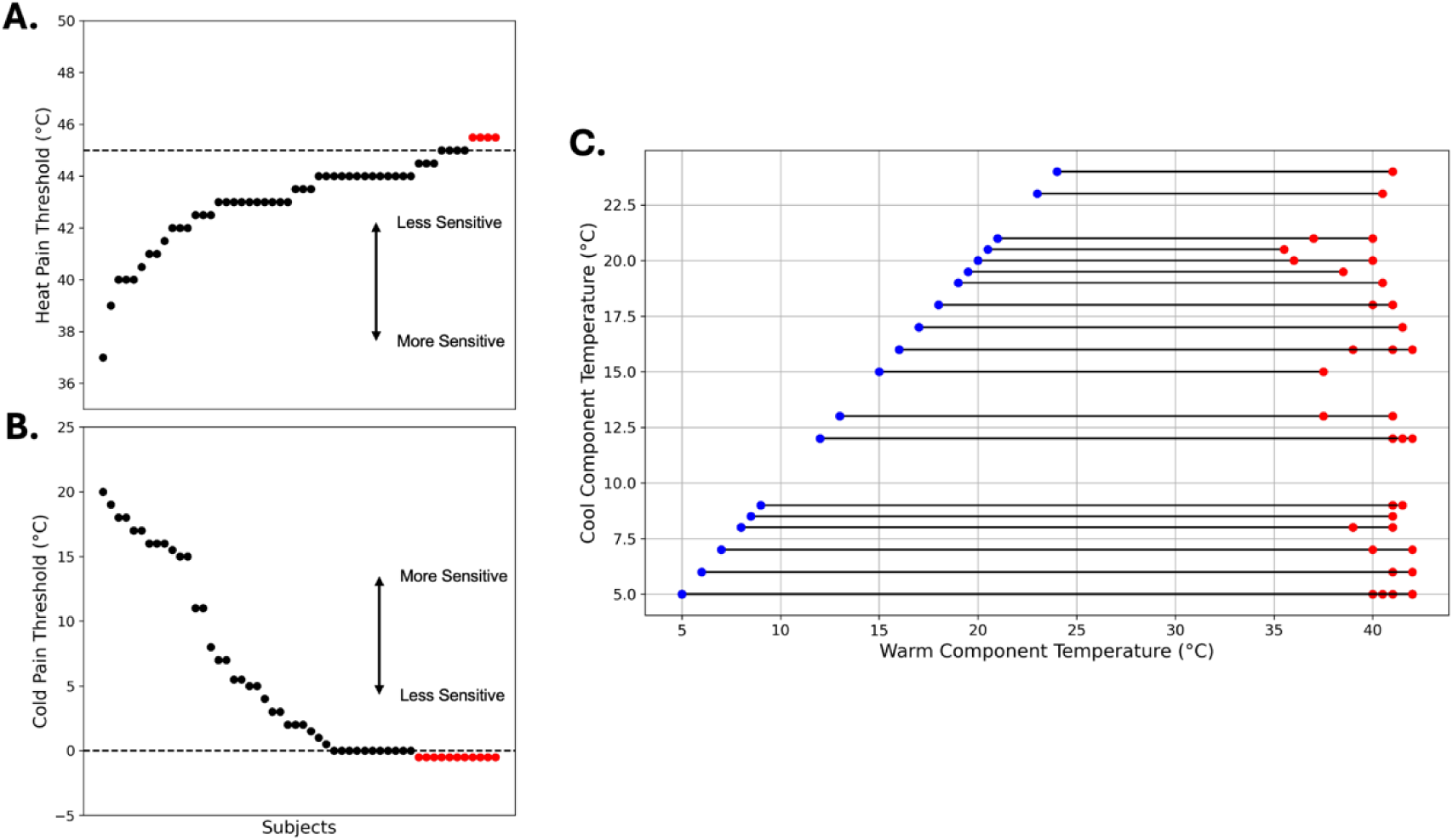
Participants’ noxious thermal stimulus thresholds and TGI stimulation temperatures. (A) Heat pain threshold and (B) cold pain thresholds for each participant. Red dots indicate participants whose HPT or CPT were outside the range of stimulation temperatures, indicated by the dotted lines (maximum 45°C and minimum 0°C). (C) TGI component temperatures and differences: blue dots represent the cool component temperature, red dots represent the warm component temperature, and the lines connecting them represent the magnitudes of difference of component temperatures. Dotted lines indicate noxious thresholds for both the heat and cold directions. Note: in panel C participants who had the same WCT and CCT combinations will have overlapping data points within the figure.

Participants received an individually calibrated thermal grill stimulus. The median CCT was 8.0°C [5.0-16.3]; males had a median CCT of 10.5°C [5.0-15.0], and females had a median of 6°C [5.0-17.5], with no significant difference between groups (p=0.780). Participants had a median WCT of 41.0°C [40.0 – 42.0]; males had a median WCTs of 41°C [40.1 – 42.0], and females had a median of 41.0°C [40.0 – 42.0], with no significant difference between groups (p = 0.662). With individual calibration of TGI component temperatures, we observed a responder rate of 78%. Participants’ CCT and WCT are summarized in Figure 4C.

### TGI Qualitative Sensations

Participants completed the MPQ following a thermal grill stimulation, to explore the qualitative descriptions commonly used to categorize sensation evoked by the TGI. The frequencies of descriptors used are provided in Figure 5. Thermal sensory descriptors were most commonly reported, with “hot” (51.9%), “cold” (44.2%) and “cool” (34.6%), followed by other sensory descriptors such as “tingling” (25%) and “spreading” (21.2%). Evaluative terms such as “annoying” (30.7%) and “troublesome” (9.6%) were also frequently reported. Affective descriptors, including “fearful” (3.8%) or “dreadful” (1.9%), were among the least frequently selected. Participants also selected descriptors within the miscellaneous categories. The frequency of all MPQ descriptors selected is provided in Table S1.

**Figure 5:**
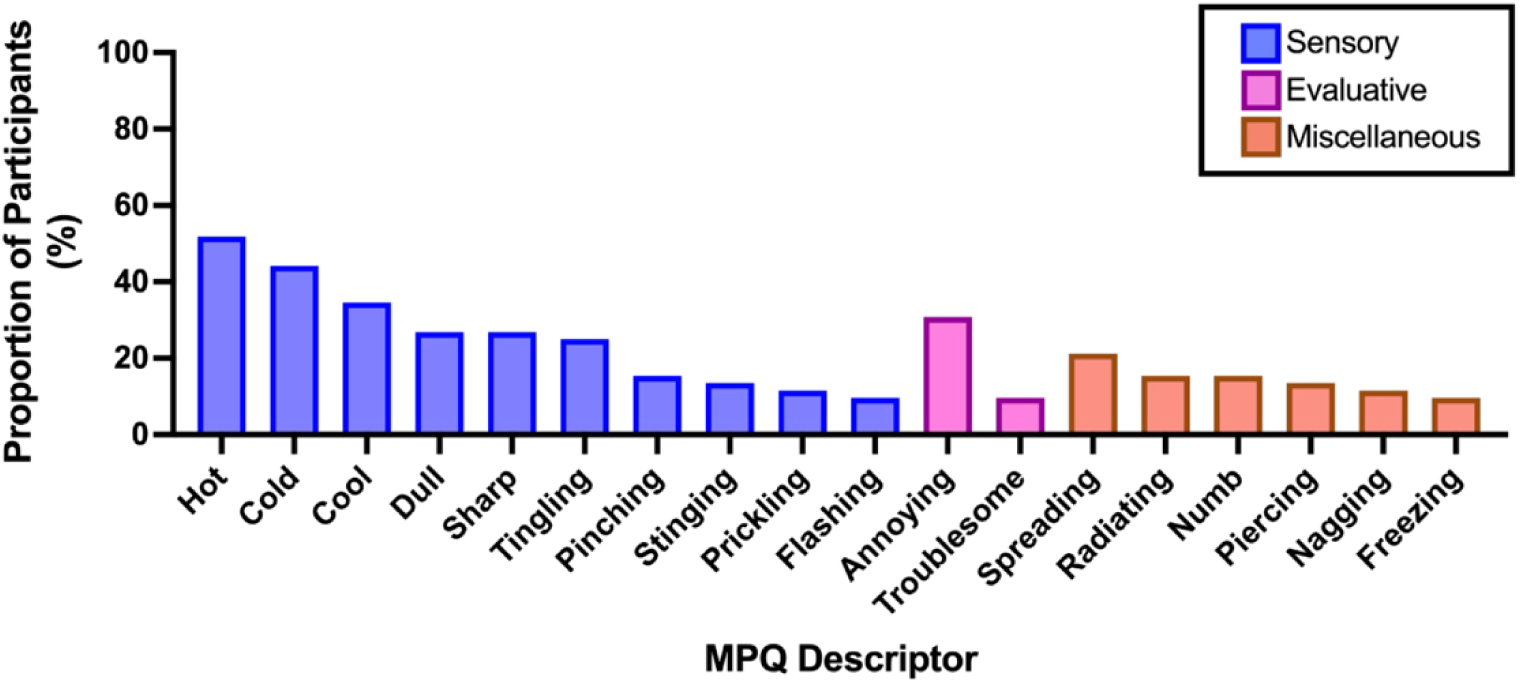
Frequency of descriptors from the McGill Pain Questionnaire used to describe the Thermal Grill Illusion in Responders. Percent frequencies of the most used descriptors (>5 participants or ∼10% of sample) from the McGill Pain Questionnaire (MPQ) used to describe the thermal grill illusion. Descriptors are categorized as sensory (blue), evaluative (green), miscellaneous (purple), or affective (of which none met the cutoff) dimensions.

### Brush Allodynia

No participants indicated any pain sensations to an innocuous brush stimulus within the stimulus site post-phTG stimulation, therefore no participant developed brush allodynia.

### Primary Hyperalgesia

Both pre-phTG and post-phTG measures were not normally distributed (both p < 0.001), thus non-parametric tests were used. Pre-phTG average pain intensity ratings ranged from 0.3 to 56.7, with a median rating [IQR range] of 9.2 [2.23 – 18.30]. Post-phTG average pain intensity ratings ranged from 0 to 80, with a median rating of 14.2 [5.00 – 30.43]. Post-phTG pain intensity ratings were significantly higher than pre-phTG (z = −3.729, p < 0.001, one-tailed, rank-biserial correlation effect size *r* = −0.517; Figure 6).

**Figure 6:**
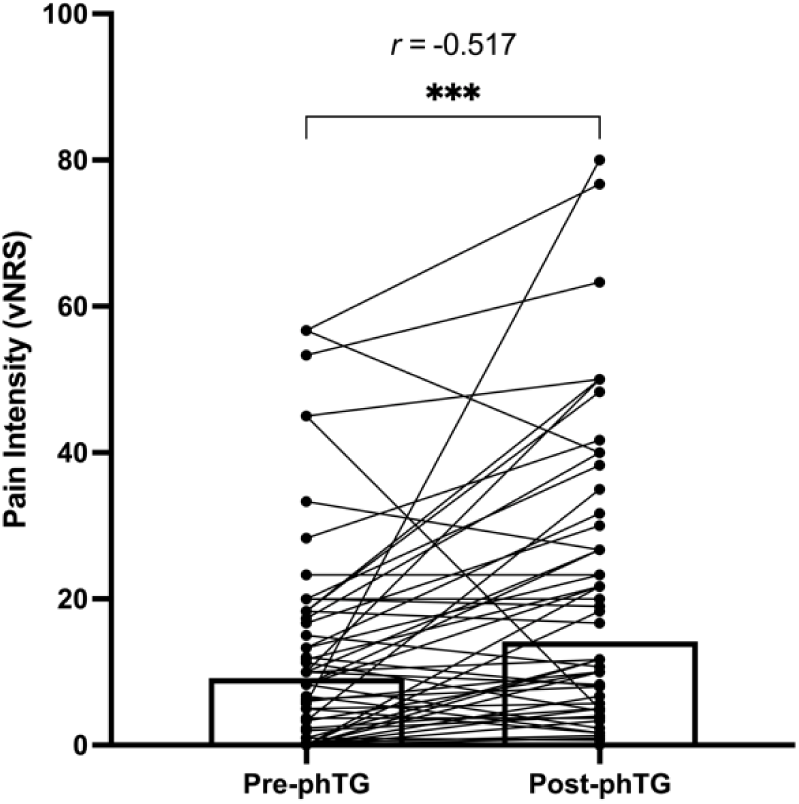
Phasic thermal grill elicits primary hyperalgesia. Pain intensity to a 512mN Pinprick before phasic thermal grill (pre-phTG) and after phTG (post-phTG) within the area of stimulation. Bars indicate median values, and each pair of dots connected by a line represents a participant. Abbreviations: r = rank-biserial correlation effect size, vNRS = verbal numerical rating scale, *** = p < 0.001.

To determine evidence for the null or alternative hypotheses, a Bayesian Wilcoxon signed rank test was performed. The analysis provided extreme evidence in favour of the alternative hypothesis (BF_+0_ = 865.185; R_hat_ = 1.004), indicating that post-phTG pain intensity was higher than pre-phTG pain intensity ratings.

### Secondary Hyperalgesia

51 out of 52 participants exhibited secondary hyperalgesia—where participants reported an area of increased pain outside of the stimulation site (area of sensitivity) which was greater than 0 cm^2^. The area of secondary hyperalgesia was normally distributed (p=0.081), thus parametric analyses were run. Participants’ calculated area of sensitivity ranged from 0 (no secondary hyperalgesia) to 93.7 cm^2^ (mean ±SD = 47.17 ±25.38 cm^2^). The area of secondary hyperalgesia evoked by phTG deviated significantly from null hypothesis of 0 cm^2^ (t = 13.404, p<0.001, one-tailed; Cohen’s *d* = 1.859; Figure 7). A Bayesian one-sample t-test revealed that there is extreme evidence in favour of the alternative hypothesis (BF_+0_ = 4.474×10^15^; error % = 4.941×10^-19^), indicating the thermal grill does elicit secondary hyperalgesia.

**Figure 7:**
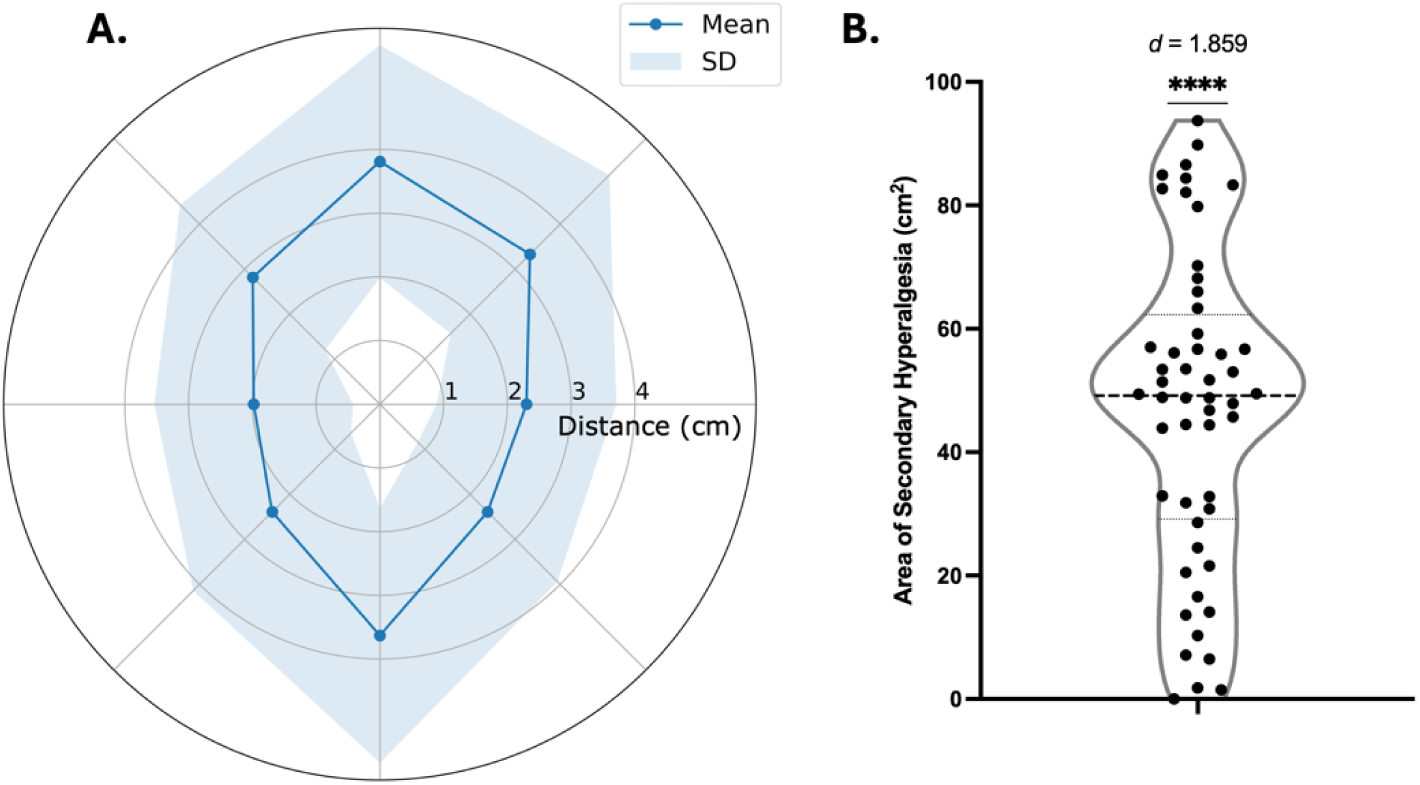
The thermal grill elicits secondary hyperalgesia. (A) The mean (blue line, shaded area: SD) area of secondary hyperalgesia elicited by the thermal grill illusion. (B) A one-tailed t-test indicates that there is a statistically significant area of secondary hyperalgesia (cm2) elicited by the thermal grill in all participants, with group median in thick dashed line and quartiles in a thinner dotted line. Abbreviations: d = Cohen’s d, SD = standard deviation, ****=p<0.0001.

### Contribution of Component Temperatures to TGI-Induced Sensitization

Despite all participants reporting both the WCT and CCT as not painful, it is still plausible that component temperatures fall within noxious ranges. Therefore, we sought to determine whether component temperatures were associated with primary and secondary hyperalgesia. A Bayesian repeated measures ANOVA with WCT and CCT as covariates was performed to determine whether the difference in mechanical pinprick pain ratings pre-versus post-phTG was influenced by component temperatures. The factor of time (pre-phTG vs post-phTG) had the highest posterior probability (P(M∣data) = 34.5%), compared to the model that included WCT (P(M∣data) = 18.8%), CCT (P(M∣data) = 29.3%), or both (P(M∣data) = 14.2%). Bayes factors (BFs) further confirmed weaker evidence for models including WCT (BF_10_ = 0.544), CCT (BF_10_ = 0.848), or both (BF_10_=0.411), compared to the model with only the factor of time (BF_10_ = 1) indicating component temperatures does not explain the significant difference between post-phTG and pre-phTG, indicating that component temperatures are not associated with primary hyperalgesia (Table S2).

A Bayesian ANCOVA for area of secondary hyperalgesia showed the null model (without WCT and CCT) had the highest posterior probability (P(M∣data) = 55.5%), compared to the model including WCT (P(M∣data) = 18.2%), CCT (P(M∣data) = 18.6%), or both (P(M∣data) = 7.6%). BFs further confirmed weaker evidence for models including WCT (BF_10_ = 0.329), CCT (BF_10_ = 0.336), or both (BF_10_ = 0.138), compared to the null model (BF_10_ = 1) that WCT or CCT does not explain the observed effect of secondary hyperalgesia following phTG (Table S2). To further explore the role of potential noxious component temperature inputs on primary outcome measures, analyses for primary and secondary hyperalgesia were rerun for all participants whose WCT and CCT were within the innocuous range (CCT > 19° and WCT < 41°C; n = 7). All primary outcome measures were normally distributed (p > 0.05), thus parametric tests were used.

For participants with innocuous component temperature inputs, there was with no significant increase in mechanical pain ratings post-phTG compared to pre-phTG (t = −1.684, p = 0.072, one-tailed, *d* = −0.637l; Figure 8A). A Bayesian paired t-test, showed anecdotal evidence in favour of the null hypothesis, indicating the thermal grill did not elicit primary mechanical hyperalgesia (BF_-0_ = 1.761; error% = 8.697 x 10^-6^). There was a significant greater area of secondary hyperalgesia than 0cm^2^ (t = 4.185, p = 0.003, one-tailed, *d* = 1.582; Figure 8B). A Bayesian one-sample t-test revealed that there is strong evidence in favour of the thermal grill eliciting secondary hyperalgesia (BF_+0_ = 20.422; error% = 8.372x 10^-7^).

**Figure 8:**
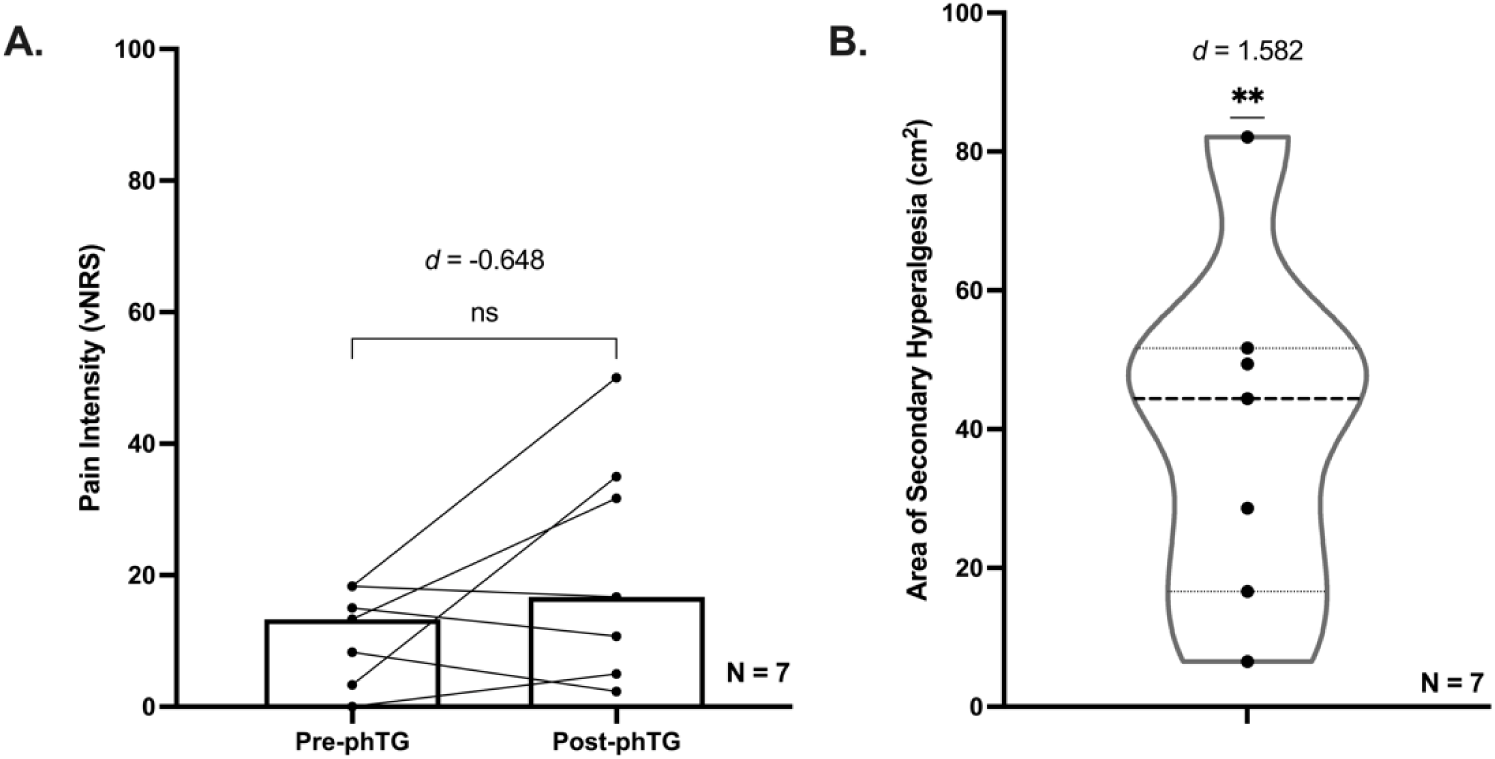
Innocuous component temperatures elicit secondary, but not primary hyperalgesia. (A) Primary hyperalgesia measures comparing mechanical punctate pain intensity pre- and post-phTG within the area of stimulation in participants with innocuous component temperatures (WCT <41°C and CCT >19°C), with median values highlighted with the bars. (B) Violin plot of area of secondary hyperalgesia (cm2) in participants with innocuous component temperatures, with group median in thick dashed line and quartiles in a thinner dotted line. Abbreviations: d = Cohen’s d, **=p<0.01.

### Sex Disaggregated and Sex Differences Analyses

Females showed no significant differences between pre- and post-phTG pain ratings (z = −1.83, p = 0.034, one-tailed, *r* = −0.346; Figure 9A), and thus did not exhibit primary hyperalgesia. Bayesian analyses showed anecdotal evidence for this observation (BF_-0_ = 1.62; R_hat_ = 1.001). In contrast, males showed a significant increase in pain ratings post-phTG (z = −3.33, p < 0.001, one-tailed, *r* = −0.653; Figure 9A), exhibiting primary hyperalgesia. Bayesian analyses further indicate extreme evidence in favour of the alternative hypothesis (BF_-0_ = 427.57; R_hat_ = 1.003). However, there was no significant difference in magnitude of primary hyperalgesia between sexes (U = 411.0, z = 1.34, p = 0.18; two-tailed, r = 0.185), with Bayesian analyses showing anecdotal-to-moderate evidence in favour of the null hypothesis of no difference between sexes (BF_10_ = 0.33; R_hat_ = 1).

**Figure 9:**
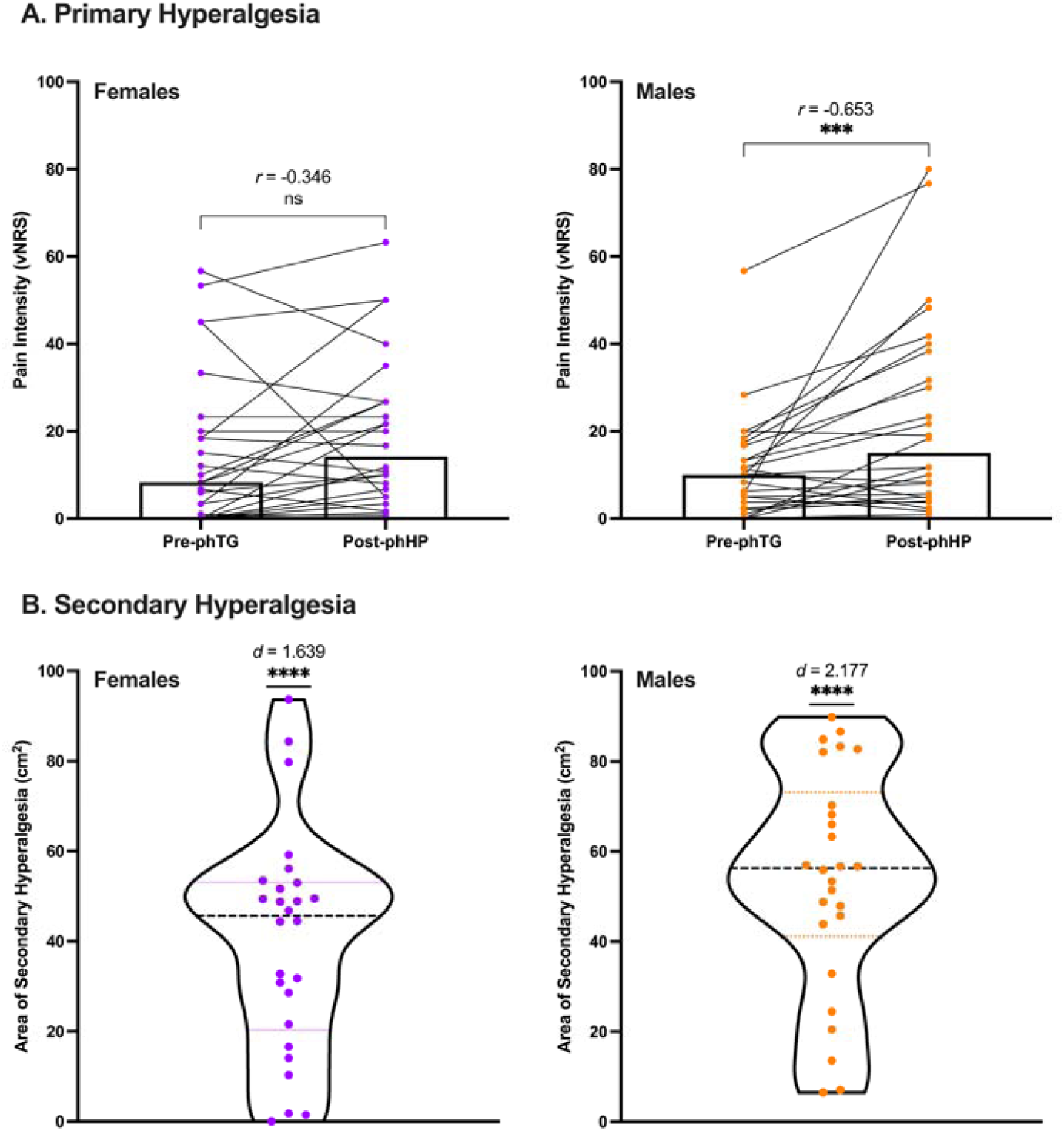
Sex disaggregated analyses of TGI-induced primary and secondary hyperalgesia. (A) Female (purple) and male (orange) primary hyperalgesia measures pre- and post-phTG. Bars represent median values. (B) Females (purple) and males (orange) area of secondary hyperalgesia (cm2), with group median in thick dashed line and quartiles in a thinner dotted line (B). Abbreviations: d = Cohen’s d, r = rank-biserial correlation effect size, ***=p<0.001, vNRS= verbal numerical pain score, ****=p<0.0001.

Both female (t = 8.36, p < 0.001, one-tailed, *d* = 1.639; Figure 9B) and male (t = 11.11, p < 0.001, one-tailed, *d* = 2.179; Figure 9B) participants showed a significant deviation in area of secondary hyperalgesia from the null hypothesis. Bayesian analyses showed extreme evidence in favour of the alternative hypothesis that both males (BF_+0_ = 5.05 x 10^8^; error% = 6.07 x 10^-12^) and females (BF_+0_ = 2.47 x 10^6^; error% = 4.26×10^-9^) developed secondary hyperalgesia. When comparing areas of secondary hyperalgesia between sexes, there was a trend towards males having larger areas, but this observation was not significant (t = −1.94, p = 0.058, two-tailed, *d* = −0.538). However, there exists anecdotal evidence in favour of the null hypothesis (BF_10_ = 0.88; error% = 0.008).

To ensure no physiological or psychosocial factor was driving any differences between sexes, independent sample t tests (parametric) and Mann Whitney U (non-parametric) were run comparing male and females’ HPT, CPT, WCT, CCT, average pain intensity during phTG, STAI State, STAI Trait, PCS, FPQ-III, and SSAS measures. There were no significant differences in any of the measures between the sexes (p > 0.05) (Table 3). Bayesian analyses supported this result finding weak evidence (1 < BF_10_ < 0.3) in favour of the null hypothesis—there are no significant differences between the groups (Table 3).

**Table 3:** Sex differences in physiological and psychosocial measures. Frequentist and complementary Bayesian analyses comparing male and female participants on physiological (HPT, CPT, WCT, CCT, and phTG pain intensity) and psychosocial (STAI State, STAI Trait, PCS, FPQ, and SSAS) measures. *Non-parametric statistics were run if either values for males or females were not normally distributed (Shapiro-Wilk test p<0.05). ** Error% (for independent samples t tests) and R_hat_ (for Mann-Whitney U tests) are provided as metrics of Bayesian model fit. Values closer to 1 for Error% and closer to 0 for R_hat_ indicate better fit and convergence, respectively. Abbreviations: MW = Mann-Whitney U Test, T = Independent samples t test.

| Measure | Normal Distribution? | Statistical test* | Frequentist Analyses |  | Bayesian Analyses |  |  |
| --- | --- | --- | --- | --- | --- | --- | --- |
| | | | Statistic | p-value | Statistic | BF $\square\square$ | Error/$R_{\text{hat}}$ ** |
| HPT | No | MW | 285 | 0.331 | 285 | 0.33 | 1 |
| CPT | No | MW | 306.5 | 0.567 | 306.5 | 0.303 | 1 |
| WCT | No | MW | 348.5 | 0.851 | 348.5 | 0.292 | 1 |
| CCT | No | MW | 306.5 | 0.559 | 306.5 | 0.308 | 1 |
| phTG Pain Intensity | No | MW | 355.5 | 0.756 | 355.5 | 0.301 | 1 |
| STAI State | No | MW | 378.5 | 0.462 | 378.5 | 0.316 | 1 |
| STAI Trait | No | MW | 373.5 | 0.521 | 373.5 | 0.303 | 1 |
| PCS | No | MW | 259.5 | 0.421 | 259.5 | 0.339 | 1 |
| FPQ | Yes | T | 1.309 | 0.197 | - | 0.562 | 0.008 |
| SSAS | Yes | T | 0.632 | 0.53 | - | 0.328 | 0.008 |

### Psychosocial Influences on TGI Sensitization

To explore whether thermal grill sensitization is associated to psychosocial measures, Bayesian correlations were performed between outcome measures—primary and secondary hyperalgesia, and TGI pain intensity during phTG—and questionnaire measures (BDI-II, STAI Trait, STAI State, PCS, FPQ-III, SSAS). There existed moderate evidence in favour of the null hypothesis— i.e., that psychosocial measures were not correlated with outcome measures (3 < BF_01_ < 6) (Figure S1).

## Discussion

The TGI, where interlaced innocuous warm and cool stimuli elicit a painful sensation, can reveal the neural underpinnings of thermonociceptive integration and perception. The neural mechanisms of the TGI remain unknown. This study explored whether the TGI elicits peripheral and/or central sensitization, as measured by brush allodynia, primary hyperalgesia and secondary hyperalgesia—each with clearly established mechanisms [77,100]. Specifically, if the TGI elicits primary hyperalgesia this indicates that the innocuous thermal inputs of the illusion activate peripheral nociceptors [57,100]; if the TGI elicits brush allodynia, then we can infer that WDR cells are likely involved [77]; and, lastly, if the TGI elicits secondary hyperalgesia, but not primary hyperalgesia or brush allodynia, then this suggests some form of integration occurs centrally at the level of the dorsal horn likely in HPC cells [30,31]— though these cells have yet to be identified in humans, and have only been characterized in non-human mammals [29,30,32,110]. We report that a phasic calibrated TGI elicits primary and secondary hyperalgesia but not brush allodynia. Furthermore, when inputs were firmly in the innocuous range (WCT <41°C and CCT >19°C), only secondary hyperalgesia was observed. These data suggest that calibrated TGI may not require integration by WDR cells but instead may be integrated in the dorsal horn by HPC cells. Primary hyperalgesia was significantly greater in males than females, suggesting sex differences in the mechanisms of integration for calibrated TGI: in males, TGI integration may occur in the periphery to some degree, whereas in females, integration may occur in the dorsal horn of the spinal cord, or supraspinally. Finally, we explored the contribution of psychosocial factors known to affect pain sensitivity to TGI sensitization measures and report no statistical relationships.

Two prominent theories about how the TGI elicits pain, the addition [16,48,49] and disinhibition theories [31,33], posit that the TGI is mediated by C-fibres with integration occurring centrally [87]; however, the C-fibres type involved remains unknown. The addition theory posits that the second-order neurons mediating the TGI are WDR [16,48,49], whereas the disinhibition theory posits HPC neurons are responsible [28,31–33].

The thermal grill, classically consisting of 40°C and 20°C, is largely thought to be innocuous [87]. The perception of pain when thermal component inputs are combined, and the absence of pain when isolated, is hallmark to the illusory phenomenon. We show the experience produces paradoxical burning and cold percepts, consistent with previous findings [7,16,53]. Further, the percepts evoked are typically sensory or evaluative, but not affective, in dimension. The paradoxical perception of pain with a fixed combination of warm and cool thermal inputs is not present in all individuals—only between 20-75% of people find the illusion painful (termed “TGI responders”) [3,16,58,80,81]. Consistent with several recent studies, we show that individually calibrated component temperatures can greatly improve TGI responsiveness (78%) [2,3,37,39,61,72].

Humans’ heat pain thresholds typically vary between 39° and 45°C, and even outside that range in some individuals [36,47,51,65,107]. The transduction of noxious heat stimuli is mediated by TRPV1 (activation threshold: 42°C), a class of temperature-sensitive receptor channels that mediate thermal encoding [56,73]. Cold pain remains less understood, and thresholds are thought to be between 5° – 20°C [67] and mediated by TRPA1 (activation threshold: 17°C) [60,62,79].

While cold pain is proposed to be mediated at least in some part by Aδ fibres [34,35,88,89], ischemic block studies have proposed a prominent role of CPNs [43,104,106]. Our observation of the abolition of primary hyperalgesia in participants with CCT > 19°C, could indicate that the group-level observations were driven by cold component temperatures that bled into the noxious range. CPNs are thought to encode noxious stimuli outside the thermoneutral zone (< 24°C) [12,18,28] and microneurographic studies have observed firing at stimuli below 19°C [19] and greater than 39°C [97]. Furthermore, evidence suggests that CPNs exhibit sensitization following mild tissue injury [97] and mild deviations outside the thermoneutral zone [12]. Therefore, there remains a possibility that the TGI does still involve some form of peripheral integration, as observed through the group-level observation of primary hyperalgesia. A potential candidate proposed to mediate the TGI peripherally are type-2 bimodal C-fibres (C2) [66], which respond to warming (>38°C) and cooling (<24°C) into the noxious ranges [18]. However, C2 fibres are mechanically insensitive and thus do not account for the primary mechanical hyperalgesia observed [18]. Therefore, the most likely candidate mediating our calibrated TGI are C-mechano-heat-cold (CMHC) fibres which begin firing <19°C [19] and this is corroborated by the absence of primary hyperalgesia in those who have a CCT >19°C.

The observation of secondary hyperalgesia, especially in the innocuous temperature subsample where primary hyperalgesia was not observed, indicates that there is some integration of the TGI within the dorsal horn. While secondary hyperalgesia is not a direct measure of central sensitization, it is a widely used behavioural correlate [64], and thus used alongside the TGI could be used to explore altered nociceptive processing [3,87,93]. Of the three major classes of second-order neurons in the dorsal horn [28], HPC and WDR neurons have been posited to mediate the TGI. NS cells have been ruled out as they exclusively respond to noxious mechanical and heat stimuli [32,109], and show no change in activity in response to thermal grill stimulation [31]. HPC neurons are insensitive to low-threshold mechanical stimulation, but show graded responses to noxious heat, mechanical, and cold stimuli [22,28–30,32,110]. Conversely, WDR neurons respond to innocuous and noxious mechanical, heat, and cold [4,24,25,45]. Both HPC and WDR neurons sensitize in response to phasic noxious stimulation or tissue injury [25,30,52,74], and are thought to contribute to secondary hyperalgesia [57,63,100]. The primary distinction between these two fibre classes is their responses to innocuous mechanical stimulation—especially in sensitized states where they mediate brush allodynia [105]. Given our observation that no participant reported pain to brush stimulation following phTG, it is TGI likely does not require WDRs, which are thought to be the mediators of stroking hyperalgesia [4,77]. Therefore, if absence of brush allodynia indicates a lack of WDR involvement, calibrated TGI is most likely integrated within the dorsal horn by HPC neurons. Tied with the observations of primary hyperalgesia with CCT <19°C, likely activating CPNs [12], and role of HPC neurons in secondary hyperalgesia, our findings are consistent with the disinhibition theory rather than the addition theory.

Secondary hyperalgesia is assumed to be elicited by tissue injury or phasic noxious stimulation (> 45°C) [75] driven by the sensitization of central nociceptors [23,86,100]. However, a few studies that have observed secondary hyperalgesia following phasic non-painful stimulation, with temperatures as low as 39 – 41°C inducing secondary hyperalgesia when applied to the skin for extended periods of time (> 15 minutes) [21]. Another study found punctate allodynia following phasic stimulation of non-painful temperatures, however some of these fell into the mild noxious range (> 42°C) [83]. Thus, our findings corroborate these data, as they demonstrate that non-painful phasic stimulation can consistently produce secondary hyperalgesia.

We also report that females did not develop primary hyperalgesia from the TGI, whereas males did. When formally comparing sexes, we did not find a statistically significant difference. As such, future studies should further investigate whether the TGI drives primary hyperalgesia between sexes, with component temperatures firmly in the innocuous ranges. Furthermore, our study did not identify any significant differences in phTG ratings, in contrast to the one study that reported that females have higher pain ratings to the TGI [6]. No other studies have investigated sex differences in TGI. We also observed anecdotal evidence in favour of males having larger areas of secondary hyperalgesia compared to females, however, this difference could be due to differences in forearm size when calculating the area of sensitization. As we did not collect forearm size, future studies are needed to parse out any sex differences in TGI sensitization, and whether mechanistic differences exist.

While we demonstrated that the TGI can elicit secondary hyperalgesia, and thus central sensitization, some limitations remain. It is unclear whether the observation of primary hyperalgesia in males resulted from component temperatures bleeding into the mild noxious range, despite non-painful reports. Given that male and female participants had similar calibrated component temperatures, further research is needed to clarify primary nociceptor activation and sensitization thresholds to mild temperatures, particularly considering sex differences. Further investigation is needed to determine if these sensitization findings can be replicated with the classic thermal grill. Additionally, our inclusion criteria (TGI pain intensity rating ≥5/100) may have biased our sample toward those sensitive to the illusion, potentially limiting generalizability.

Despite these limitations, the current study is the first to show that the TGI can induce sensitization. We used these altered states to explore the potential peripheral and spinal mechanisms of the TGI, showing that the TGI can induce central sensitization, and may be mediated by HPC cells, rather than WDRs. Given that altered thermonociceptive processing is common in chronic pain conditions [102,108] and that the TGI has been used to study these differences [3,93], our findings suggest that the TGI could be a valuable tool for investigating pathological pain states and their underlying mechanisms.

## Acknowledgements

MAC was funded by a Natural Sciences and Engineering Research Council (NSERC) Canada Graduate Scholarship – Master’s. The authors have no conflicts to report. This work was funded through MM’s discretionary funds, and a NSERC Discovery Grant (RGPIN-2018-4908 to MM). MM holds a Canada Research Chair (Tier 2) in Pain NeuroImaging and is supported by a University of Toronto Centre for the Study of Pain Scientist Award. We would like to thank Karen D. Davis for her advice and contributions to the study design.

## References

[1] Abssy SS, Osborne NR, Osokin EE, Tomin R, Honigman L, Khan JS, Vera NWD, Furman A, Mazaheri A, Seminowicz DA, Moayedi M. It’s the Sound, not the Pulse: Peripheral Magnetic Stimulation Reduces Central Sensitization through Auditory Modulatory Effects. eLife 13, 2024.

[2] Adam F, Alfonsi P, Kern D, Bouhassira D. Relationships between the paradoxical painful and nonpainful sensations induced by a thermal grill. Pain 155:2612–2617, 2014.

[3] Adam F, Jouët P, Sabaté JM, Perrot S, Franchisseur C, Attal N, Bouhassira D. Thermal grill illusion of pain in patients with chronic pain: a clinical marker of central sensitization? Pain 164:638–644, 2023.

[4] Akiyama T, Nagamine M, Davoodi A, Ivanov M, Carstens MI, Carstens E. Innocuous warming enhances peripheral serotonergic itch signaling and evokes enhanced responses in serotonin-responsive dorsal horn neurons in the mouse. J Neurophysiol 117:251–259, 2017.

[5] Alrutz S. On the temperature-senses. Mind 7:141–144, 1898.

[6] Averbeck B, Seitz L, Kolb FP, Kutz DF. Sex differences in thermal detection and thermal pain threshold and the thermal grill illusion: a psychophysical study in young volunteers. Biol Sex Differ 8:29, 2017.

[7] Bach P, Becker S, Kleinbohl D, Holzl R. The thermal grill illusion and what is painful about it. Neurosci Lett 505:31–35, 2011.

[8] Baeumler PI, Conzen P, Irnich D. High Temporal Summation of Pain Predicts Immediate Analgesic Effect of Acupuncture in Chronic Pain Patients-A Prospective Cohort Study. Front Neurosci 13:498, 2019.

[9] Barsky AJ, Wyshak G, Klerman GL. The somatosensory amplification scale and its relationship to hypochondriasis. J Psychiatr Res 24:323–334, 1990.

[10] Beck AT, Steer RA, Brown G. Beck depression inventory–II. Psychological assessment, 1996.

[11] Berry KJ, Johnston JE, Mielke PW. Ordinal-Level Variables, I. The Measurement of Association: A Permutation Statistical Approach. Cham: Springer International Publishing, 2018. pp. 223-295.

[12] Blomqvist A. Pain and temperature, and human awareness: The legacy of Bud Craig. Temperature (Austin) 10:395–401, 2023.

[13] Blomqvist A, Zhang E-T, Craig AD. Cytoarchitectonic and immunohistochemical characterization of a specific pain and temperature relay, the posterior portion of the ventral medial nucleus, in the human thalamus. Brain 123:601–619, 2000.

[14] Boettger MK, Grossmann D, Bar KJ. Thresholds and perception of cold pain, heat pain, and the thermal grill illusion in patients with major depressive disorder. Psychosom Med 75:281–287, 2013.

[15] Boettger MK, Schwier C, Bar KJ. Sad mood increases pain sensitivity upon thermal grill illusion stimulation: implications for central pain processing. Pain 152:123–130, 2011.

[16] Bouhassira D, Kern D, Rouaud J, Pelle-Lancien E, Morain F. Investigation of the paradoxical painful sensation (’illusion of pain’) produced by a thermal grill. Pain 114:160–167, 2005.

[17] Cain DM, Khasabov SG, Simone DA. Response properties of mechanoreceptors and nociceptors in mouse glabrous skin: an in vivo study. J Neurophysiol 85:1561–1574, 2001.

[18] Campero M, Baumann TK, Bostock H, Ochoa JL. Human cutaneous C fibres activated by cooling, heating and menthol. J Physiol 587:5633–5652, 2009.

[19] Campero M, Serra J, Ochoa JL. C-polymodal nociceptors activated by noxious low temperature in human skin. JPhysiol(Lond) 497:565–572, 1996.

[20] Caterina MJ, Schumacher MA, Tominaga M, Rosen TA, Levine JD, Julius D. The capsaicin receptor: a heat-activated ion channel in the pain pathway. Nature 389:816–824, 1997.

[21] Cervero F, Gilbert R, Hammond RGE, Tanner J. Development of secondary hyperalgesia following non-painful thermal stimulation of the skin: a psychophysical study in man. Pain 54:181–189, 1993.

[22] Christensen BN, Perl ER. Spinal neurons specifically excited by noxious or thermal stimuli: marginal zone of the dorsal horn. J Neurophysiol 33:293–307, 1970.

[23] Coderre TJ, Melzack R. Cutaneous hyperalgesia: contributions of the peripheral and central nervous systems to the increase in pain sensitivity after injury. Brain Res 404:95–106, 1987.

[24] Coghill RC. The Distributed Nociceptive System: A Framework for Understanding Pain. Trends Neurosci 43:780–794, 2020.

[25] Coghill RC, Mayer DJ, Price DD. Wide dynamic range but not nociceptive-specific neurons encode multidimensional features of prolonged repetitive heat pain. J Neurophysiol 69:703–716, 1993.

[26] Cohen J. Statistical power analysis for the behavioral sciences. Hillside, NJ: Lawrence Erlbaum Associates, 1988.

[27] Cohen J. A power primer. Psychol Bull 112:155–159, 1992.

[28] Craig AD. Pain mechanisms: labeled lines versus convergence in central processing. Annu Rev Neurosci 26:1–30, 2003.

[29] Craig AD. Lamina I, but not lamina V, spinothalamic neurons exhibit responses that correspond with burning pain. Journal of Neurophysiology 92:2604–2609, 2004.

[30] Craig AD, Andrew D. Responses of spinothalamic lamina I neurons to repeated brief contact heat stimulation in the cat. Journal of Neurophysiology 87:1902–1914, 2002.

[31] Craig AD, Bushnell MC. The Thermal Grill Illusion: Unmasking the Burn of Cold Pain. Science 265:252–255, 1994.

[32] Craig AD, Krout K, Andrew D. Quantitative response characteristics of thermoreceptive and nociceptive lamina I spinothalamic neurons in the cat. Journal of Neurophysiology 86:1459–1480, 2001.

[33] Craig AD, Reiman EM, Evans A, Bushnell MC. Functional imaging of an illusion of pain. Nature 384:258–260, 1996.

[34] Davis KD. Cold-induced pain and prickle in the glabrous and hairy skin. Pain 75:47–57, 1998.

[35] Davis KD, Dostrovsky JO. Perception of cold pain in the glabrous and hairy skin of normal subjects: contribution of A- and C-fiber primary afferent nociceptors. Am Pain Soc J, 1994.

[36] Defrin R, Shachal-Shiffer M, Hadgadg M, Peretz C. Quantitative somatosensory testing of warm and heat-pain thresholds: the effect of body region and testing method. Clin J Pain 22:130–136, 2006.

[37] Deolindo CS, Ehmsen JF, Courtin AS, Mitchell AG, Kraenge CE, Nikolova N, Allen MG, Fardo F. Assessing individual sensitivity to the Thermal Grill Illusion: A two-dimensional adaptive psychophysical approach. J Pain 27:104732, 2024.

[38] Fardo F, Beck B, Allen M, Finnerup NB. Beyond labeled lines: A population coding account of the thermal grill illusion. Neurosci Biobehav Rev 108:472–479, 2020.

[39] Fardo F, Finnerup NB, Haggard P. Organization of the Thermal Grill Illusion by Spinal Segments. Ann Neurol 84:463–472, 2018.

[40] Ferre ER, Iannetti GD, van Dijk JA, Haggard P. Ineffectiveness of tactile gating shows cortical basis of nociceptive signaling in the Thermal Grill Illusion. Sci Rep 8:6584, 2018.

[41] Folstein MF, Folstein SE, McHugh PR. “Mini-mental state”. A practical method for grading the cognitive state of patients for the clinician. J Psychiatr Res 12:189–198, 1975.

[42] Forstenpointner J, Berry D, Baron R, Borsook D. The cornucopia of central disinhibition pain - An evaluation of past and novel concepts. Neurobiol Dis 145:105041, 2020.

[43] Fruhstorfer H. Thermal sensibility changes during ischemic nerve block. Pain 20:355–361, 1984.

[44] Fruhstorfer H, Harju E-L, Lindblom UF. The significance of A-δ and C fibres for the perception of synthetic heat. European Journal of Pain 7:63–71, 2003.

[45] Giere C, Melchior M, Dufour A, Poisbeau P. Spinal integration of hot and cold nociceptive stimuli by wide-dynamic-range neurons in anesthetized adult rats. Pain Rep 6:e983, 2021.

[46] Goss-Sampson M. Statistical analysis in JASP: A guide for students: JASP, 2019.

[47] Granovsky Y, Matre D, Sokolik A, Lorenz J, Casey KL. Thermoreceptive innervation of human glabrous and hairy skin: a contact heat evoked potential analysis. PAIN 115:238–247, 2005.

[48] Green BG. Synthetic heat at mild temperatures. Somatosens Mot Res 19:130–138, 2002.

[49] Green BG. Temperature perception and nociception. J Neurobiol 61:13–29, 2004.

[50] Green BG, Akirav C. Threshold and rate sensitivity of low-threshold thermal nociception. Eur J Neurosci 31:1637–1645, 2010.

[51] Greene LC, Hardy JD. Spatial summation of pain. J Appl Physiol 13:457–464, 1958.

[52] Guan Y, Raja SN. Wide-dynamic-range neurons are heterogeneous in windup responsiveness to changes in stimulus intensity and isoflurane anesthesia level in mice. Journal of Neuroscience Research 88:2272–2283, 2010.

[53] Harper DE, Hollins M. Coolness both underlies and protects against the painfulness of the thermal grill illusion. Pain 155:801–807, 2014.

[54] Hermesdorf M, Berger K, Baune BT, Wellmann J, Ruscheweyh R, Wersching H. Pain Sensitivity in Patients With Major Depression: Differential Effect of Pain Sensitivity Measures, Somatic Cofactors, and Disease Characteristics. J Pain 17:606–616, 2016.

[55] Hillary FG, Medaglia JD. What the replication crisis means for intervention science. Int J Psychophysiol 154:3–5, 2020.

[56] Hoffstaetter LJ, Bagriantsev SN, Gracheva EO. TRPs, et al.: a molecular toolkit for thermosensory adaptations. Pflugers Arch 470:745–759, 2018.

[57] Hsieh MT, Donaldson LF, Lumb BM. Differential contributions of A- and C-nociceptors to primary and secondary inflammatory hypersensitivity in the rat. Pain 156:1074–1083, 2015.

[58] Hunter J, Dranga R, van Wyk M, Dostrovsky JO. Unique influence of stimulus duration and stimulation site (glabrous vs. hairy skin) on the thermal grill-induced percept. Eur J Pain 19:202–215, 2015.

[59] Inoue Y, Gerrett N, Ichinose-Kuwahara T, Umino Y, Kiuchi S, Amano T, Ueda H, Havenith G, Kondo N. Sex differences in age-related changes on peripheral warm and cold innocuous thermal sensitivity. Physiol Behav 164:86–92, 2016.

[60] Jürgens TP, Sawatzki A, Henrich F, Magerl W, May A. An improved model of heat-induced hyperalgesia--repetitive phasic heat pain causing primary hyperalgesia to heat and secondary hyperalgesia to pinprick and light touch. PLoS One 9:e99507, 2014.

[61] Karashima Y, Talavera K, Everaerts W, Janssens A, Kwan KY, Vennekens R, Nilius B, Voets T. TRPA1 acts as a cold sensor in vitro and in vivo. Proc Natl Acad Sci U S A 106:1273–1278, 2009.

[62] Kern D, Pelle-Lancien E, Luce V, Bouhassira D. Pharmacological dissection of the paradoxical pain induced by a thermal grill. Pain 135:291–299, 2008.

[63] Kwan KY, Corey DP. Burning cold: involvement of TRPA1 in noxious cold sensation. J Gen Physiol 133:251–256, 2009.

[64] LaMotte RH, Shain CN, Simone DA, Tsai EF. Neurogenic hyperalgesia: psychophysical studies of underlying mechanisms. J Neurophysiol 66:190–211, 1991.

[65] Leone CM, Lenoir C, van den Broeke EN. Assessing signs of central sensitization: A critical review of physiological measures in experimentally induced secondary hyperalgesia. Eur J Pain, 2024.

[66] Li X, Petrini L, Defrin R, Madeleine P, Arendt-Nielsen L. High resolution topographical mapping of warm and cold sensitivities. Clin Neurophysiol 119:2641–2646, 2008.

[67] Lloyd-Smith DL, Mendelssohn K. Tolerance limits to radiant heat. Br Med J 1:975–978, 1948.

[68] Ma Q. Labeled lines meet and talk: population coding of somatic sensations. J Clin Invest 120:3773–3778, 2010.

[69] Magerl W, Krumova EK, Baron R, Tolle T, Treede RD, Maier C. Reference data for quantitative sensory testing (QST): refined stratification for age and a novel method for statistical comparison of group data. Pain 151:598–605, 2010.

[70] Marotta A, Ferre ER, Haggard P. Transforming the thermal grill effect by crossing the fingers. Curr Biol 25:1069–1073, 2015.

[71] McNeil DW, Rainwater AJ, 3rd. Development of the Fear of Pain Questionnaire--III. J Behav Med 21:389–410, 1998.

[72] Melzack R. The McGill Pain Questionnaire: major properties and scoring methods. Pain 1:277–299, 1975.

[73] Michaelides A, Zis P. Depression, anxiety and acute pain: links and management challenges. Postgrad Med 131:438–444, 2019.

[74] Mitchell AG, Ehmsen JF, Christensen DE, Stuckert AV, Haggard P, Fardo F. Disentangling the spinal mechanisms of illusory heat and burning sensations in the thermal grill illusion. Pain 165:2370–2378, 2024.

[75] Paricio-Montesinos R, Schwaller F, Udhayachandran A, Rau F, Walcher J, Evangelista R, Vriens J, Voets T, Poulet JFA, Lewin GR. The Sensory Coding of Warm Perception. Neuron 106:830–841 e833, 2020.

[76] Pertovaara A. A neuronal correlate of secondary hyperalgesia in the rat spinal dorsal horn is submodality selective and facilitated by supraspinal influence. Exp Neurol 149:193–202, 1998.

[77] Quesada C, Kostenko A, Ho I, Leone C, Nochi Z, Stouffs A, Wittayer M, Caspani O, Brix Finnerup N, Mouraux A, Pickering G, Tracey I, Truini A, Treede RD, Garcia-Larrea L. Human surrogate models of central sensitization: A critical review and practical guide. Eur J Pain 25:1389–1428, 2021.

[78] Raphael KG, Marbach JJ, Gallagher RM. Somatosensory amplification and affective inhibition are elevated in myofascial face pain. Pain Med 1:247–253, 2000.

[79] Sadeghi M, Manaheji H, Zaringhalam J, Haghparast A, Nazemi S, Bahari Z, Noorbakhsh SM. Evaluation of the GABAA Receptor Expression and the Effects of Muscimol on the Activity of Wide Dynamic Range Neurons Following Chronic Constriction Injury of Sciatic Nerve in Rats. Basic Clin Neurosci 12:651–666, 2021.

[80] Salomons TV, Moayedi M, Erpelding N, Davis KD. A brief cognitive-behavioural intervention for pain reduces secondary hyperalgesia. Pain 155:1446–1452, 2014.

[81] Sawada Y, Hosokawa H, Hori A, Matsumura K, Kobayashi S. Cold sensitivity of recombinant TRPA1 channels. Brain Res 1160:39–46, 2007.

[82] Schaldemose EL, Horjales-Araujo E, Svensson P, Finnerup NB. Altered thermal grill response and paradoxical heat sensations after topical capsaicin application. Pain 156:1101–1111, 2015.

[83] Schaldemose EL, Raaschou-Nielsen L, Bohme RA, Finnerup NB, Fardo F. It is one or the other: No overlap between healthy individuals perceiving thermal grill illusion or paradoxical heat sensation. Neurosci Lett 802:137169, 2023.

[84] Scheuren R, Sutterlin S, Anton F. Rumination and interoceptive accuracy predict the occurrence of the thermal grill illusion of pain. BMC Psychol 2:22, 2014.

[85] Schifftner C, Schulteis G, Wallace MS. Effect of Intravenous Alfentanil on Nonpainful Thermally Induced Hyperalgesia in Healthy Volunteers. J Clin Pharmacol 57:1207–1214, 2017.

[86] Schneider CA, Rasband WS, Eliceiri KW. NIH Image to ImageJ: 25 years of image analysis. Nature methods 9:671–675, 2012.

[87] Schreiber S, Hewitt D, Seymour B, Yoshida W. Enhancing experimental design through Bayes factor design analysis: insights from multi-armed bandit tasks [version 1; peer review: 3 approved with reservations]. Wellcome Open Research 9, 2024.

[88] Schulte H, Sollevi A, Segerdahl M. The distribution of hyperaemia induced by skin burn injury is not correlated with the development of secondary punctate hyperalgesia. J Pain 5:212–217, 2004.

[89] Shin DA, Chang MC. A Review on Various Topics on the Thermal Grill Illusion. J Clin Med 10, 2021.

[90] Simone DA, Kajander KC. A-delta nociceptors are excited by noxious cold stimuli. SocNeurosciAbstr 19:324, 1993.

[91] Simone DA, Kajander KC. Responses of cutaneous A-fiber nociceptors to noxious cold. Journal of Neurophysiology 77:2049–2060, 1997.

[92] Spielberger CD. State-trait anxiety inventory for adults. 1983.

[93] Story GM, Peier AM, Reeve AJ, Eid SR, Mosbacher J, Hricik TR, Earley TJ, Hergarden AC, Andersson DA, Hwang SW, McIntyre P, Jegla T, Bevan S, Patapoutian A. ANKTM1, a TRP-like channel expressed in nociceptive neurons, is activated by cold temperatures. Cell 112:819–829, 2003.

[94] Sullivan MJL, Bishop SR, Pivik J. The Pain Catastrophizing Scale: Development and validation. Psychological Assessment 7:524–532, 1995.

[95] Sumracki NM, Buisman-Pijlman FT, Hutchinson MR, Gentgall M, Rolan P. Reduced response to the thermal grill illusion in chronic pain patients. Pain Med 15:647–660, 2014.

[96] Thunberg T. Förnimmelserna vid till samma ställe lokaliserad, samtidigt pågående köld-och värmeretning. Uppsala Läkfören Förh 2:489–495, 1896.

[97] Thunberg T, Harper D. Co-localized sensations resulting from simultaneous cold and warm stimulation (Förnimmelserne vid till samma ställe lokaliserad, samtidigt pågående köld-och värmeretning), 2016.

[98] Tillman DB, Treede RD, Meyer RA, Campbell JN. Response of C fibre nociceptors in the anaesthetized monkey to heat stimuli: correlation with pain threshold in humans. J Physiol 485 (Pt 3):767–774, 1995.

[99] Torebjork E. Nociceptor activation and pain. Philos Trans R Soc Lond B Biol Sci 308:227–234, 1985.

[100] Treede RD, Meyer RA, Campbell JN. Comparison of heat and mechanical receptive fields of cutaneous C-fiber nociceptors in monkey. J Neurophysiol 64:1502–1513, 1990.

[101] Treede RD, Meyer RA, Campbell JN. Myelinated mechanically insensitive afferents from monkey hairy skin: heat-response properties. J Neurophysiol 80:1082–1093, 1998.

[102] Treede RD, Meyer RA, Raja SN, Campbell JN. Peripheral and central mechanisms of cutaneous hyperalgesia. Prog Neurobiol 38:397–421, 1992.

[103] van Doorn J, van den Bergh D, Bohm U, Dablander F, Derks K, Draws T, Etz A, Evans NJ, Gronau QF, Haaf JM, Hinne M, Kucharsky S, Ly A, Marsman M, Matzke D, Gupta A, Sarafoglou A, Stefan A, Voelkel JG, Wagenmakers EJ. The JASP guidelines for conducting and reporting a Bayesian analysis. Psychon Bull Rev 28:813–826, 2021.

[104] Vierck CJ, Wong F, King CD, Mauderli AP, Schmidt S, Riley JL, 3rd. Characteristics of sensitization associated with chronic pain conditions. Clin J Pain 30:119–128, 2014.

[105] Wagenmakers EJ, Marsman M, Jamil T, Ly A, Verhagen J, Love J, Selker R, Gronau QF, Smira M, Epskamp S, Matzke D, Rouder JN, Morey RD. Bayesian inference for psychology. Part I: Theoretical advantages and practical ramifications. Psychon Bull Rev 25:35–57, 2018.

[106] Wahren LK, Torebjork E, Jorum E. Central suppression of cold-induced C fibre pain by myelinated fibre input. Pain 38:313–319, 1989.

[107] Witting N, Kupers RC, Svensson P, Arendt-Nielsen L, Gjedde A, Jensen TS. Experimental brush-evoked allodynia activates posterior parietal cortex. Neurology 57:1817–1824, 2001.

[108] Yarnitsky D, Ochoa JL. Release of cold-induced burning pain by block of cold-specific afferent input. Brain 113:893–902, 1990.

[109] Yarnitsky D, Sprecher E, Zaslansky R, Hemli JA. Heat pain thresholds: normative data and repeatability. Pain 60:329–332, 1995.

[110] Zeng X, Sun Y, Zhiying Z, Hua L, Yuan Z. Chronic pain-induced functional and structural alterations in the brain: A multi-modal meta-analysis. J Pain:104740, 2025.

[111] Zhang TC, Janik JJ, Grill WM. Mechanisms and models of spinal cord stimulation for the treatment of neuropathic pain. Brain Res 1569:19–31, 2014.

[112] Zhang X, Davidson S, Giesler GJ, Jr. Thermally identified subgroups of marginal zone neurons project to distinct regions of the ventral posterior lateral nucleus in rats. J Neurosci 26:5215–5223, 2006.

[71] Vierck CJ, Wong F, King CD, Mauderli AP, Schmidt S, Riley JL, 3rd. Characteristics of sensitization associated with chronic pain conditions. Clin J Pain 30:119–128, 2014.

[72] Wagenmakers EJ, Marsman M, Jamil T, Ly A, Verhagen J, Love J, Selker R, Gronau QF, Smira M, Epskamp S, Matzke D, Rouder JN, Morey RD. Bayesian inference for psychology. Part I: Theoretical advantages and practical ramifications. Psychon Bull Rev 25:35–57, 2018.

[73] Wahren LK, Torebjork E, Jorum E. Central suppression of cold-induced C fibre pain by myelinated fibre input. Pain 38:313–319, 1989.

[74] Witting N, Kupers RC, Svensson P, Arendt-Nielsen L, Gjedde A, Jensen TS. Experimental brush-evoked allodynia activates posterior parietal cortex. Neurology 57:1817–1824, 2001.

[75] Yarnitsky D, Ochoa JL. Release of cold-induced burning pain by block of cold-specific afferent input. Brain 113:893–902, 1990.

[76] Zeng X, Sun Y, Zhiying Z, Hua L, Yuan Z. Chronic pain-induced functional and structural alterations in the brain: A multi-modal meta-analysis. J Pain:104740, 2025.

[77] Zhang X, Davidson S, Giesler GJ, Jr. Thermally identified subgroups of marginal zone neurons project to distinct regions of the ventral posterior lateral nucleus in rats. J Neurosci 26:5215–5223, 2006.

