## Supplementary material for "The Thermal Grill Elicits Central Sensitization": Figure S1 (from another version of this preprint: version 2, 2025-02-15)

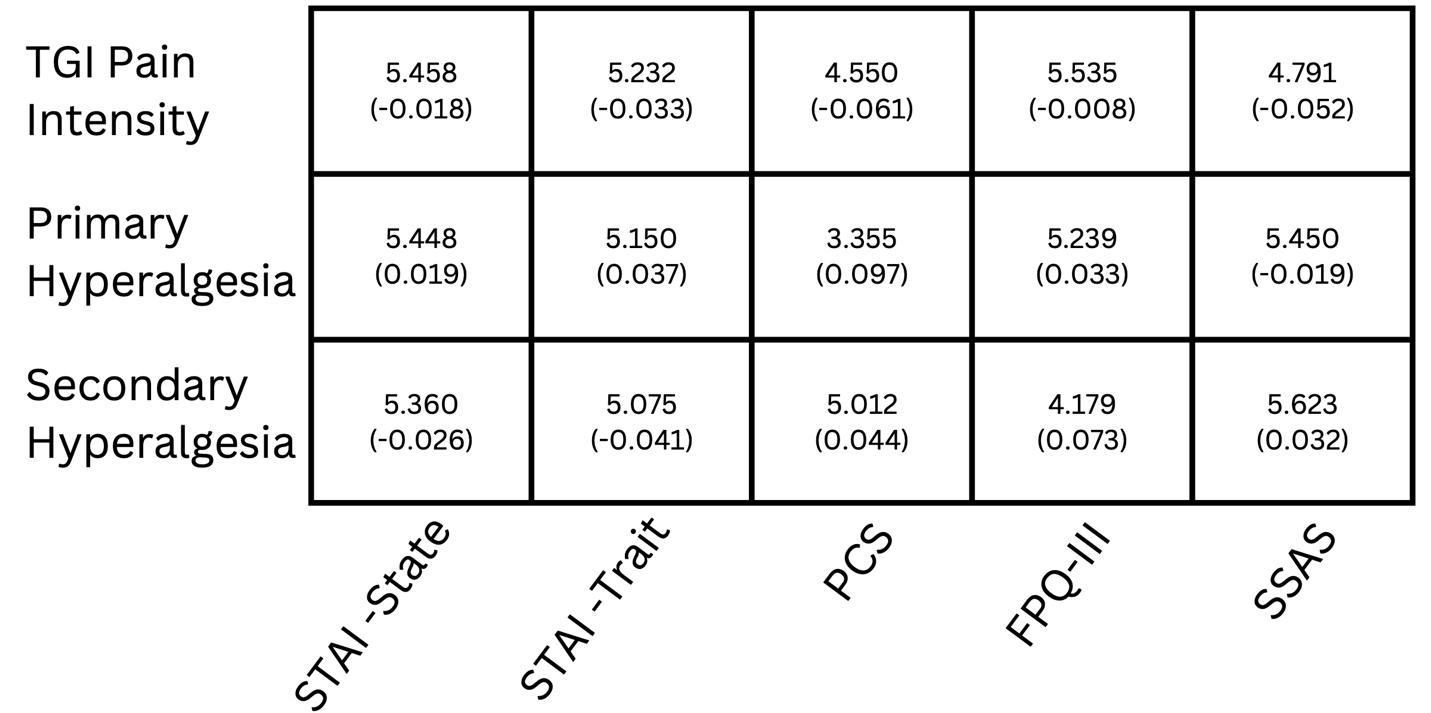

**Supplementary Figure 1**: **Correlation matrix between psychosocial questionnaire measures and primary outcome measures**. Correlations between psychosocial measures and three outcome measures (primary and secondary hyperalgesia, and average TGI pain intensity rating during phTG). Bayes Factor (BF_01_) with Kendall’s tau correlation values in parentheses.
