## Supplementary material for "The Thermal Grill Elicits Central Sensitization": Table S1 (from another version of this preprint: version 2, 2025-02-15)

**Supplementary Table 1:** **McGill Pain Questionnaire descriptor frequencies.** Participants were prompted to select from a list of descriptive words on the MPQ to describe the sensations evoked by the TG stimulation. Frequency and percentage frequency of words selected are summarized.

| **MPQ Word** | **Frequency Count** | **Frequency Percentage (%)** |
| --- | --- | --- |
| *Hot* | 27 | 51.9% |
| *Cold* | 23 | 44.2% |
| *Cool* | 18 | 34.6% |
| *Annoying* | 16 | 30.8% |
| *Dull* | 14 | 26.9% |
| *Sharp* | 14 | 26.9% |
| *Tingling* | 13 | 25.0% |
| *Spreading* | 11 | 21.2% |
| *Radiating* | 8 | 15.4% |
| *Pinching* | 8 | 15.4% |
| *Numb* | 8 | 15.4% |
| *Piercing* | 7 | 13.5% |
| *Stinging* | 7 | 13.5% |
| *Nagging* | 6 | 11.5% |
| *Prickling* | 6 | 11.5% |
| *Troublesome* | 5 | 9.6% |
| *Freezing* | 5 | 9.6% |
| *Flashing* | 5 | 9.6% |
| *Pressing* | 4 | 7.7% |
| *Heavy* | 4 | 7.7% |
| *Intense* | 4 | 7.7% |
| *Boring* | 4 | 7.7% |
| *Tender* | 4 | 7.7% |
| *Burning* | 4 | 7.7% |
| *Pulsing* | 3 | 5.8% |
| *Aching* | 3 | 5.8% |
| *Stabbing* | 3 | 5.8% |
| *Cramping* | 3 | 5.8% |
| *Sore* | 3 | 5.8% |
| *Flickering* | 3 | 5.8% |
| *Splitting* | 2 | 3.8% |
| *Searing* | 2 | 3.8% |
| *Rasping* | 2 | 3.8% |
| *Fearful* | 2 | 3.8% |
| *Tiring* | 2 | 3.8% |
| *Hurting* | 2 | 3.8% |
| *Pulling* | 2 | 3.8% |
| *Quivering* | 2 | 3.8% |
| *Tugging* | 2 | 3.8% |
| *Cutting* | 2 | 3.8% |
| *Tight* | 2 | 3.8% |
| *Punishing* | 2 | 3.8% |
| *Taut* | 2 | 3.8% |
| *Nauseating* | 1 | 1.9% |
| *Drawing* | 1 | 1.9% |
| *Pricking* | 1 | 1.9% |
| *Shooting* | 1 | 1.9% |
| *Squeezing* | 1 | 1.9% |
| *Beating* | 1 | 1.9% |
| *Dreadful* | 1 | 1.9% |
| *Gnawing* | 1 | 1.9% |
| *Wretched* | 1 | 1.9% |
| *Pounding* | 1 | 1.9% |
| *Penetrating* | 1 | 1.9% |
| *Scalding* | 1 | 1.9% |
| *Smarting* | 1 | 1.9% |
| *Itchy* | 1 | 1.9% |
| *Wretching* | 1 | 1.9% |
