## Supplementary material for "The Thermal Grill Elicits Central Sensitization": Table S2 (from another version of this preprint: version 2, 2025-02-15)

**Supplementary Table 2:** **Bayesian ANCOVA analyses exploring the contribution of component temperatures on outcome measures.** (A) Bayesian ANCOVA analysis to determine whether WCT or CCT have significant contributions to secondary hyperalgesia measures. (B) Bayesian repeated measures ANCOVA to determine whether WCT or CCT have significant contributions to changes in pain intensity ratings to mechanical pinprick stimulations post- compared to pre-phTG.

| *A. Secondary Hyperalgesia Bayesian ANCOVA* | | |  |  |  |  |
| --- | --- | --- | --- | --- | --- | --- |
| **Models** | **P(M)** | **P(M\|data)** | **BF_M_** | **BF_10_** | **error %** | **BF_incl_*** |
| Null model | 0.25 | 0.555 | 3.736 | 1 |  |  |
| CCT | 0.25 | 0.186 | 0.687 | 0.336 | 0.002 | 0.357 |
| WCT | 0.25 | 0.182 | 0.67 | 0.329 | 0.002 | 0.349 |
| WCT + CCT | 0.25 | 0.076 | 0.248 | 0.138 | 0.011 |  |
| *B. Primary Hyperalgesia Bayesian Repeated Measures ANCOVA* | | | | |  |  |
| **Models** | **P(M)** | **P(M\|data)** | **BF_M_** | **BF_10_** | **error %** |  |
| Timepoint | 0.125 | 0.345 | 3.695 | 1 |  | 30.831 |
| Timepoint + CCT | 0.125 | 0.293 | 2.901 | 0.848 | 2.46 |  |
| Timepoint + WCT | 0.125 | 0.188 | 1.621 | 0.544 | 2.233 |  |
| Timepoint + WCT + CCT | 0.125 | 0.142 | 1.16 | 0.411 | 3.047 |  |
| Null model (incl. subject and random slopes) | 0.125 | 0.012 | 0.082 | 0.034 | 1.906 |  |
| CCT | 0.125 | 0.01 | 0.068 | 0.028 | 2.261 | 0.815 |
| WCT | 0.125 | 0.006 | 0.042 | 0.017 | 2.274 | 0.516 |
| WCT + CCT | 0.125 | 0.004 | 0.029 | 0.012 | 1.979 |  |

*BF_incl_ = Bayes factor for average of effects across models. Note that 0<BF<3 indicate anecdotal evidence for H_1_; 3<BF<10 indicates moderate evidence for H_1_; 10<BF<30 indicates strong evidence for H_1_, and 30<BF<100 indicates very strong evidence for H_1_; 100<BF indicates extreme evidence for H_1_.
